# Mitochondrial oxidation of the carbohydrate fuel driven by pyruvate dehydrogenase robustly enhances stemness of older and geriatric Intestinal Stem Cells

**DOI:** 10.1101/2024.09.23.614374

**Authors:** Syed Ahmed, Aasem Awwad, Nerise Eddy, Garrett Weber, Zrar Shahid, Zubin Sethi, Jonathan Labampa, Robert Murphy, Eric W. Roth, Kyle Gustafson, Hardik Shah, Sinju Sundaresan

## Abstract

**Background and Aims:** Aging impairs Intestinal Stem Cell (ISC) function and attenuates their regenerative capacity. Although the transcriptional landscape governing ISC fate during aging has been described, almost nothing is known about how metabolite handling regulates ISC renewal and maintains stemness. We investigated how mitochondrial metabolism of glucose and fatty acid-derived carbons, regulated by the gatekeeper, pyruvate dehydrogenase (PDH) rescues ISC stemness in older and geriatric mice and humans.

**Methods:** Proximal small intestinal organoids (enteroids) generated from pinch biopsy specimens obtained from young (21-25y) and older individuals (64-75y), and GFP-sorted single ISCs from Lgr5-EGFP mice (2-24 months) were used to examine hallmarks of ISC stemness. Mitochondrial morphology was evaluated using transmission electron microscopy. Mitochondrial oxygen consumption rate (OCR), ATP (mitoATP), and glycolytic ATP production were measured in the presence of full and single metabolic substrates (pyruvate, glutamate, and fatty acids) in whole cell and isolated mitochondria using the high throughput Seahorse XF technology. Carbon flux through TCA cycle was determined by ^13^C_6_-glucose tracing and measuring ^13^C enrichment in TCA cycle intermediates using liquid chromatography mass spectrometry.

**Results:** Age induced decline in ISC stemness is driven by a dramatic decrease in PDH activity that shuttles pyruvate away from the TCA cycle. Restoring PDH activity by inhibition of pyruvate dehydrogenase kinase 4 (PDK4) drives glucose-derived carbon entry into TCA cycle and subsequently increases mitochondrial OCR and mitoATP, collectively rescuing the decline in stemness in aging ISCs. The observed shift in fuel preference from fatty acids to glucose is unaltered by PDK4 inhibition.

**Conclusion:** PDH upregulation rescues age-induced decline in ISC stemness in humans and mice via directing glucose derived carbons to TCA cycle and increasing mitoATP production.

## INTRODUCTION

Aging impairs Intestinal Stem Cell (ISC) function and attenuates their regenerative capacity, resulting in decreased nutrient absorption, reduced barrier function, increased susceptibility to infections and bleeding, and slow recovery from radiation damage ^1^. Impaired ISC function also constitutes one of the underlying mechanisms for age-associated intestinal disorders and intestinal cancer due to altered functional competition among aged ISCs. Molecular mechanisms regulating aging of mammalian ISCs are only beginning to be unraveled ^2^. Although the transcriptional landscape governing ISC function has been studied, very little is known about how metabolite handling regulates ISC fate and renewal. Nutrient-derived metabolites induce chromatin reshaping, epigenetic modifications, and modulate gene expression ^3, 4^, yet signaling targets and molecular machinery underlying ISC fate and function are largely unknown.

The pyruvate dehydrogenase complex (PDC) that catalyzes the oxidative decarboxylation of pyruvate to acetyl-CoA, acts as a gatekeeper in metabolism by linking the glycolytic pathway to mitochondrial oxidative phosphorylation (OXPHOS) ^5–7^. The PDCs in prokaryotes and eukaryotes are composed of three enzymes: pyruvate dehydrogenase (PDH), dihydrolipoamide acetyltransferase, and dihydrolipoamide dehydrogenase which sequentially catalyze the oxidative decarboxylation of pyruvate ^8, 9^. The flux of pyruvate through PDC is tightly regulated by two regulatory enzymes, pyruvate dehydrogenase kinase (PDK) ^10, 11^ and pyruvate dehydrogenase phosphatase (PDP) ^12–14^, which regulate PDC activity by phosphorylation (inhibition) and dephosphorylation (activation) of serine residues 232, 293, and 300 of PDH. PDKs regulate the metabolic switch between glycolysis and OXPHOS and have been reported to play crucial roles in tumor cell proliferation, migration, invasion, and apoptosis resistance ^9^.

We report, for the first time, novel and significant roles of PDK4 and mitochondrial OXPHOS in regulation of ISC fate in mouse and human intestines. Using human duodenal enteroid lines and Lgr5-EGFP mice we demonstrate that aging-induced decline in ISC stemness is associated with a shift in preference from fatty acids (FAs) to glucose as metabolic fuel source. Metabolite flux analyses using liquid chromatography-mass spectrometry on GFP^high^ Lgr5^+^ ISCs demonstrate progressive increase in anaerobic pyruvate utilization, along with a concomitant decrease in FA utilization with age. Seahorse-based bioenergetics analyses revealed a corresponding increase in mitochondrial oxygen consumption rate (OCR) and ATP (mitoATP) production. Mechanistically, dramatic decrease in PDH activity during aging shuttles pyruvate away from the tricarboxylic acid (TCA) cycle. Restoring PDH activity by PDK4 inhibition drives glucose and glucose-derived carbon entry into TCA cycle, subsequently increasing mitochondrial OCR and mitoATP, collectively reversing the decline in stemness in aging ISCs. Our data demonstrate novel and robust therapeutic potential of the PDH-PDK4 axis in rescuing age-induced decline in ISC stemness and lays mechanistic foundations for future intervention studies.

## METHODS

### Mice

All experiments were approved by Midwestern University’s Institutional Committee on the Use and Care of Animals. Male and female *Lgr5-EGFP-IRES-CreERT2* (*Lgr5,* Jackson Labs, #008875) mice ^15^ were housed in the Midwestern University vivarium at 23°C on a 0600–1800-h light cycle, with *ad libitum* access to standard chow and water.

### Human and mouse enteroid culture

Human duodenal enteroid lines were generated from pinch biopsy specimens as described ^16^. Enteroids from mice were harvested as described ^16–18^ with modifications. Refer to Supplemental Methods for details.

### Transfection of human and GFP^high^ mouse ISCs

Human duodenal enteroid suspensions and sorted GFP^high^ mouse ISCs were transfected using Lipofectamine 2000, as described ^19^ with modifications. After 2 hours of plating, siRNA (5nM-10nM) and lipofectamine complexes were added, as per manufacturer’s instructions (Invitrogen, Inc). Lipofectamine and siRNA (Origene) were added separately to DMEM media with 10% dialyzed FBS (1:1), incubated (5 min), and then 500Lμl of formed siRNA complex medium was bathed over matrigel. Addition of serum robustly enhanced internalization of siRNA into the Matrigel and enteroids ^19^ performed as indicated. After 48 hours of transfection, media was replaced, and assays were

### Isolation of human and mice ISC mitochondria

Two-day-old human duodenal enteroids and enteroids generated from GFP^high^ mouse ISCs were dislodged by gentle trituration using ice cold PBS (without Ca^2+^/Mg^2+^). Enteroid suspensions were centrifuged (350*g*, 5 min, 4°C) and pellets were solubilized in Mitochondrial Extraction Buffer (MEB; 0.25 M sucrose, 20 mM HEPES-KOH, 10 mM KCl, 1.5 mM MgCl2, 1 mM EDTA, 1 mM EGTA, 1 mM dithiothreitol, 0.1 mM PMSF). Homogenates were centrifuged (750*g*, 10 min, 4°C) and pellets were resuspended in extraction buffer, homogenized, and centrifuged as described above. The supernatants from two low-speed spins were pooled and then centrifuged (10,000*g*, 15 min, 4°C). Crude mitochondria, recovered in the pellets, were resuspended in MEB, and filtered (100 µm) for OCR analyses.

### Measurement of OCR

Measurement of OCR of enteroids with full access to all metabolic substrates was performed using a Seahorse XF24 analyzer, as per standard protocols (Agilent Technologies, Inc) with modifications ^20^. Briefly, about 17-25 enteroids resuspended in 3µL of Matrigel (1:1) were plated on the center of each well of the microplate, in between the 3 nodes. After 2 days, basal and maximal OCR was measured in the presence of the mitochondrial inhibitor oligomycin (1.2 μM), the mitochondrial uncoupler FCCP (5 μM), and the respiratory chain inhibitor rotenone (1 μM). Mitochondrial and glycolytic ATP was measured using the Real Time ATP Assay kit (Agilent Technologies, Inc). Data was normalized to DNA content.

### Measurement of OCR in isolated mitochondria

Freshly isolated mitochondria (200-300 enteroids for 10 µg of mitochondria) were plated in each well of an XF24 plate in 50 μl of MEB (with 0.1% FA–free bovine serum albumin). Plates were centrifuged (2500*g*, 4°C, 20 min at 2000*g*) and then 450 μl of MEB (5 mM pyruvate/5 mM malate, 5 mM glutamate/5 mM malate, 40 μM palmitoyl-CoA/40 μM carnitine/5 mM malate, or 10 mM succinate) was added to each well. Plates were incubated in a non-CO_2_ incubator for 8-10 min. OCRs were measured in the presence of ADP (4 mM), oligomycin (1.5 μM), FCCP (4 μM), and rotenone (1 μM). For OCR in isolated mitochondria, a shorter run time (∼40 minutes) was used to preserve viability during the assay.

### 13C6-glucose tracing in TCA cycle intermediates using liquid chromatography tandem mass spectrometry (LC-MS)

Two-three-day old enteroids (generated from GFP^high^ sorted ISCs), grown in monolayers were switched to glucose free media for 3 hours and incubated in 5mm D-glucose (unlabeled) or ^13^C_6_- glucose (Cambridge Isotope Laboratories, Inc) for 6 hours. Enteroids were extracted in dry ice-cold 80% methanol for LC-MS analyses as described ^21, 22^. MS detection was performed using Orbitrap IQ-X Tribrid mass spectrometer with H-ESI probe. Data was acquired using the Xcalibur software (70-1000 m/z, 60k resolution). Metabolite identification was done by matching the retention time and MS/MS fragmentation to reference standards. Peak integration was performed using Skyline software ^22^ and natural abundance correction was performed using IsoCor software ^21^.

### PDH and CPT1 activity, pyruvate, and acetyl-CoA measurements

Freshly isolated whole cell or mitochondrial fractions from GFP^high^ sorted mouse ISCs or human enteroid cells were lysed (50 mM tris-HCl, 1% NP-40, 150 mM NaCl, 1 mM EDTA, 1 mM NaF, 1 mM Na_3_VO_4_) for PDH activity (Sigma Aldrich, Inc). CPT1 activity was analyzed as described ^23, 24^ by measuring the release of CoA-SH from palmitoyl-CoA using 5, 5’-Dithiobis (2-nitrobenzoic acid, DNTB) based buffer (116 mM Tris-HCl, 2.5 mM EDTA, 2 mM DTNB, 0.2% Triton X-100). Pyruvate and acetyl-CoA levels were measured using commercial kits (BioVision, Inc).

### Transmission Electron Microscopy (TEM)

Enteroids were fixed in paraformaldehyde, glutaraldehyde and osmium tetroxide (1%), dehydrated in graded series of ethanol and acetone prior to infiltration and embedding in EMBed812 epoxy resin. Sections (60nm) were cut with a diamond knife on a Leica UC7 ultramicrotome and collected on slotted grids with formvar/carbon membrane prior to staining with 3% uranyl acetate in 50% methanol and Reynold’s lead citrate solutions. Grids were analyzed using JEOL JEM 1400 Flash LaB6 emission TEM. Images were obtained with Gatan OneView digital camera at 4k resolution. Mitochondrial area and feret distance were calculated using Image J.

### Mito Tracker Assay

GFP^high^ sorted ISCs were grown in monolayers in 8-well chambers coated with collagen (20μg/cm^2^). Monolayers were incubated with 50 nM Mito Tracker dye (MitoTracker^®^ Red CMXRos, 30 minutes, 37°C), fixed with ice-cold methanol (15 min, -20°C), washed, and mounted for imaging.

### Western Blot

Please refer to Supplemental Methods.

### RT-qPCR

Please refer to Supplemental Methods.

### Statistical Analyses

Please refer to Supplemental Methods.

## RESULTS

### Progressive decline in ISC stemness in aged humans and mice

ISC stemness was evaluated by determining growth and budding capacities of human enteroid lines, harvested from duodenal pinch biopsy specimens from healthy, young (21-25y) and older (64-75y) individuals. Subject demographics are presented in Supplemental Table 3. In young enteroids, buds appeared within 3 days and became mature and prolific by the end of 6 days (**Figure 1A**). Bud appearance was significantly delayed in enteroids from older individuals. No buds were observed until after 6 days in culture (**Figure 1B**) and fewer stunted buds lacking the prolific phenotype were observed after 10 days. Morphometric analyses revealed that enteroid area was 30-43% lower (**Figure 1C**) and bud numbers were 77-93% fewer (**Figure 1D**) in enteroids from older individuals. Enteroid forming efficiency (EFE), calculated as the percentage of enteroids per 10 primary enteroids, was drastically reduced in older enteroids. EFE dropped by 82% by the end of the 8^th^ passage (**Figure 1E**).

**Figure 1.**
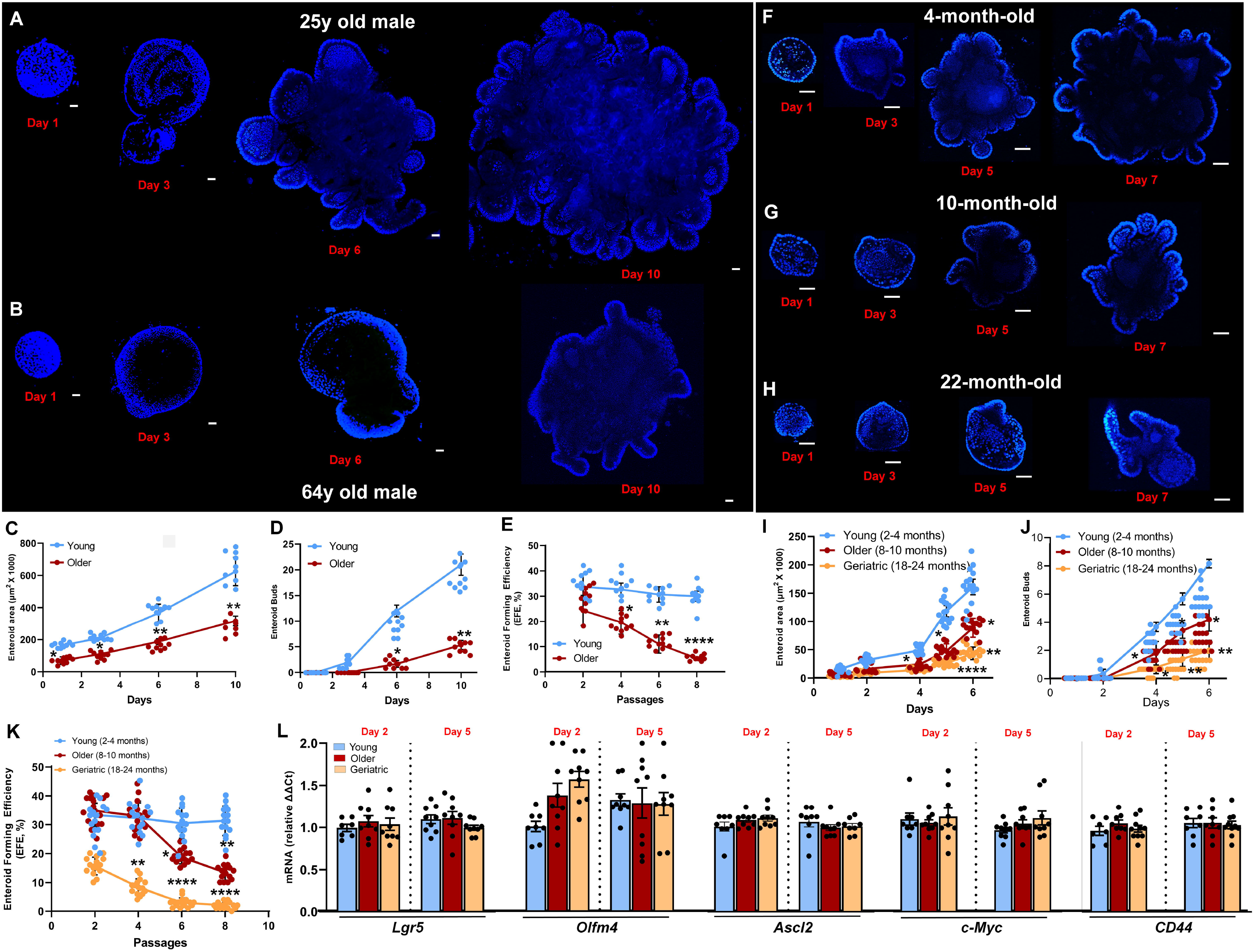
Progressive decline in ISC stemness in aged humans and mice. Representative nuclear (DAPI) staining of duodenal enteroids generated from pinch biopsy-derived enteroid lines from 25y old male **(A)** and 64y old male **(B)**. Enteroid area **(C)** and bud numbers **(D)** analyzed using linear mixed effects modeling with likelihood ratio tests (3-4 enteroids/passage/subject, 2-3 passages/subject, *n*=3 subjects/group). Enteroid Forming Efficiency (**E,** EFE, 4-6 replicates/passage/subject; *n*=3 subjects/group, 2 passages/subject). Data in (C-E) were analyzed using non-parametric repeated one-way ANOVA and post hoc Dunn’s test with Benjamini-Hochberg correction for multiple comparisons. Representative nuclear (DAPI) staining of proximal small intestinal enteroids generated from 4- **(F)**, 10- **(G)**, and 22-month-old mice **(H)**. Enteroid area **(I)** and bud numbers **(J)** analyzed as in (C-E) (3-4 enteroids/passage/mouse; *n*=5-6 mice/group). EFE (**K,** 3-4 replicates/passage/mouse; *n*=5-6 mice/group, 2 passages/mouse). Asterisks denote significance compared to the young at the given time point. *Lgr5*, *Olfm4*, *Ascl2*, *c-Myc*, *CD44* mRNA expressions **(L)** in GFP-sorted cells (GFP^high^) from proximal small intestinal crypts (∼16 cm distal to pyloric sphincter) of young, older, and geriatric Lgr5-EGFP mice and grown in culture for 2-5 days, normalized to *Gapdh,* and expressed relative to the young (*n*=mean of triplicates from 7-9 mice/age group). Heteroscedastic data was log-transformed and analyzed using repeated one-way ANOVA and post hoc Dunn’s test with Benjamini-Hochberg correction. Scale bars: (A-B, F-H) – 100 µm.

A similar pattern was observed in proximal intestinal enteroids harvested from mice (**Figure 1F-K**). Stemness declined progressively in enteroids from 2-4- (young), 8-10- (older), and 18–24-month-old (geriatric) mice. Enteroid area was reduced by 40-60% and 65-80% in older and geriatric mice, respectively (**Figure 1I**). Bud numbers followed a similar pattern (**Figure 1J**). EFE was significantly lower in geriatric mice and progressively worsened over 4 passages (**Figure 1K**). Next, we examined the impact of aging on molecular signatures of ISC stemness. We sorted for GFP-labeled ISCs from proximal small intestines of Lgr5-EGFP mice (GFP^high^) by flow cytometry and measured gene expression of key ISC markers. No significant differences were observed in *Lgr5*, *Olfm4*, *Ascl2*, *c-Myc*, and *Cd44* gene expressions (**Figure 1L**), indicating ISC function rather than absolute numbers may be impacted with aging.

### Aging reprograms ISC cellular metabolism by limiting mitochondrial utilization of FAs and mitochondrial ATP

Mitochondrial metabolism and respiration have been shown to play pivotal roles in ISC fate decisions that directly impact stemness ^25–27^. Given the decline in stemness, we first analyzed mitochondrial morphology and numbers across all age groups. Mitotracker (MT) staining was used to label mitochondria in GFP^high^ ISCs. Mitochondrion area, mean branch length, diameter, and number of branches per mitochondrion were analyzed using the Fiji Mitochondrial Analyzer plugin ^28^. We found that mitochondrion area decreases progressively with age **(Figure 2A-C, D)**. Branches of older and geriatric mitochondria were 50%-62% shorter than younger counterparts **(Figure 2A-C, E)**, possibly implying enhanced fission or reduced fusion with increasing age. No significant differences were observed in branch diameter and number of branches per mitochondrion **(Figure 2F, G)**. Ultramicroscopic evaluation of the mitochondria revealed that geriatric ISCs had significantly lower (∼40%) ferret distance compared to young ISCs **(Figures 2H-J),** implying greater sphericity. Mitochondrial numbers were comparable across all age groups (data not shown).

**Figure 2.**
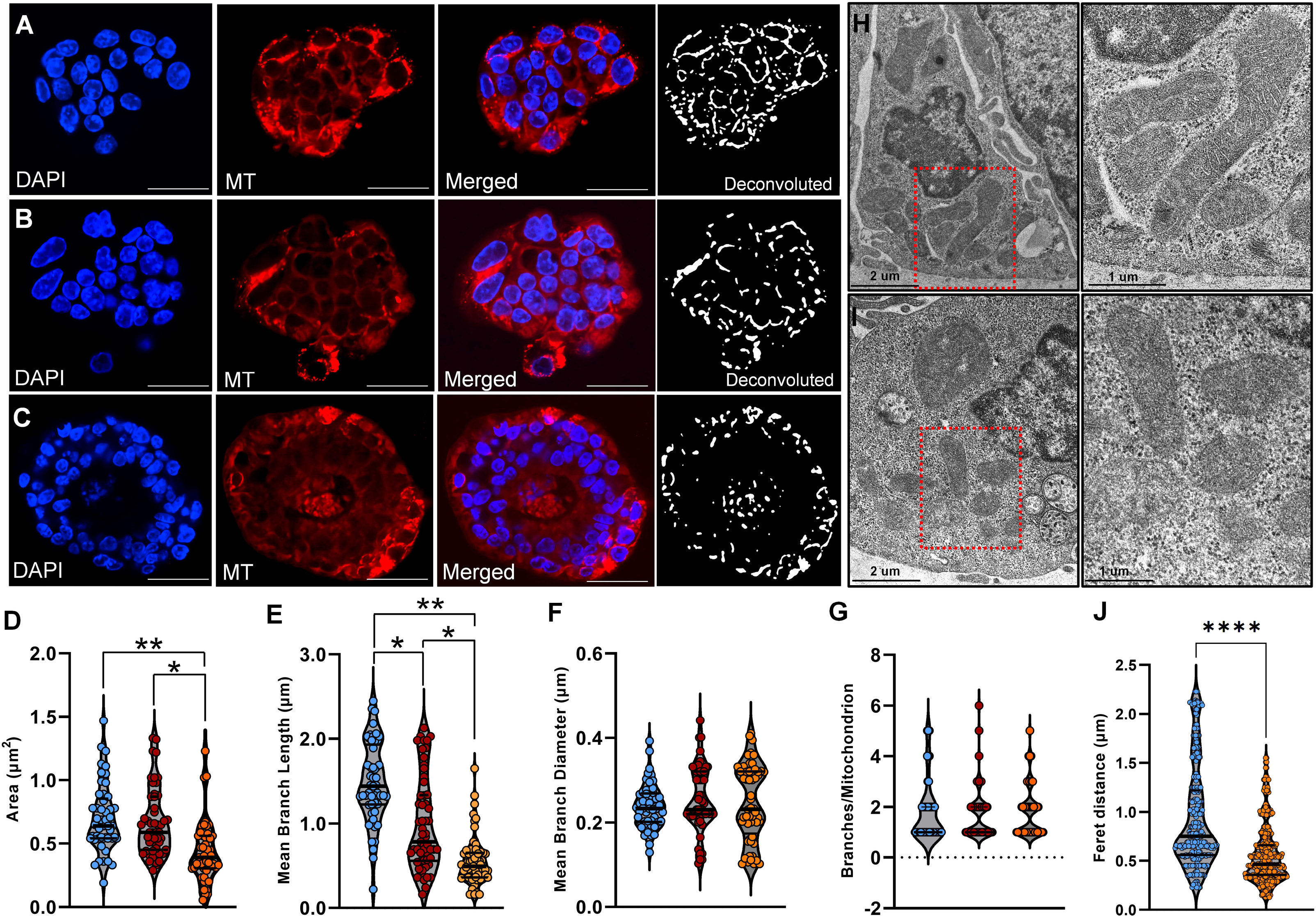
Reduced area and altered sphericity in older and geriatric ISC mitochondria. Red MitoTracker^®^ staining of enteroids harvested from 4- **(A)**, 10- **(B)**, and 22-month-old Lgr5-EGFP mice **(C)** and grown in monolayers for 2-3 days. Network representations for confocal images after deconvolution using the Fiji plugin, Mitochondrial Analyzer. Area **(D)**, mean branch length **(E)**, mean branch diameter **(F)**, and branches per ISC mitochondrion **(G)**, analyzed using Kruskal Wallis ANOVA and post hoc Dunn’s test (10-18 mitochondria/3-6 ISCs/enteroid). At least 3 enteroids were analyzed per mouse (*n*=5-6 mice/group). **(H, I)** Ultramicroscopic examination of ISC mitochondria in enteroids harvested from proximal small intestines of 4- **(H)** and 22-month-old mice **(I).** Ferret distance in ISC mitochondria **(J)** measured using Image J (12- 16 mitochondria/3-5 ISCs/enteroid; 3-4 enteroids/mouse; *n*=3 mice/group), analyzed using Wilcoxon Rank Sum test (*, p<0.05; **, p<0.01; ****, p<0.0001). Scale bars: (A-C) – 20 µm, (H, I) - 2 µm, insets - 1 µm.

To determine whether aging alters mitochondrial metabolite handling, we measured gene expression of carriers for uptake of long chain FAs (*Cpt1a* and *Cpt2*), and β-oxidation, or FA oxidation (FAO) enzymes (*Acaa2*, *Acsl1*, *Hadh*) in GFP^high^ ISCs. In older and geriatric ISCs, gene expressions of carriers and all FAO enzymes were significantly downregulated **(Supplemental Figure 1A**), while expression of rate-limiting glycolytic enzymes, phosphofructokinase (*Pfk*) and pyruvate kinase (*Pk2*) were significantly upregulated **(Supplemental Figure 1B),** suggesting that glycolysis may be augmented upon aging. To test whether aging influences FAO dependance for stemness, we evaluated enteroid budding capacities after treatment with etomoxir, the carnitine palmitoyl transferase I (CPT1) inhibitor that blocks mitochondrial FAO transport. Etomoxir (50 µM) markedly reduced growth and budding capacities of young enteroids **(Supplemental Figure 2A-C)**. Acetate, by providing an alternative source of acetyl-CoA rescued etomoxir-mediated FAO inhibition **(Supplemental Figure 2A-C)**. Suppression of growth and budding capacities, and acetate rescue were less pronounced and virtually absent in older **(Supplemental Figure 2D-F)** and geriatric enteroids **(Supplemental Figure 2G-I)**, respectively. These data suggest that FAO may be the less preferred route for ATP production in ISCs upon aging. We then determined whether propensity towards reduced FAO is due to possible differences in CPT1 activity. Indeed, CPT1 activity declined progressively with age, with dramatic decrease observed in geriatric ISCs (∼95% lower, **Supplemental Figure 3A).** This suggests that FAs may be less preferred metabolic substrate in aging ISCs, consequently implying an increased dependance on glucose for ATP.

To delineate the route for ATP production, we first measured mitochondrial OCR in GFP^high^ ISCs sorted from Lgr5-EGFP mice using Sea-Horse. When provided full access to all metabolic substrates, geriatric ISCs showed significantly lower basal (B) and maximal (M) OCRs, compared to the young (**Figure 3A, B**). Basal OCRs in geriatric ISCs were 55.4% and 44.8% lower than younger and older ISCs, respectively (**Figure 3B**, *left panel*). Maximal OCR declined progressively with age (older- 43.5% lower, geriatric-60.8% lower; **Figure 3B**, *left panel*). Similarly, basal and maximal OCRs in enteroids from older individuals were 43% and 52% lower than those of younger enteroids, respectively (**Figure 3B, *right panel***). We then determined whether altered OCR impacted ATP production by measuring mito- and glyco-ATP production rates using the real time ATP assay. As shown in **Figure 3C**, basal mitoATP decreased progressively with age, corresponding to decreased OCRs. MitoATP in geriatric ISCs was 52% lower than that of young. In contrast, glycoATP increased slightly in older and geriatric ISCs compared to the young, conceivably due to the diversion of pyruvate from mitochondrial oxidation to lactate in the cytosol. Determination of extracellular acidification rates (ECAR) that measures lactate production confirmed these findings. Geriatric ISCs had higher ECARs than young and older ISCs, at baseline and in response to mitochondrial inhibition by oligomycin (**Supplemental Figure 3B**). Similarly, enteroids from older individuals had 53.5% lower basal mitoATP while basal glycoATP remained significantly elevated (84.7% higher; **Figure 3D**). In response to oligomycin inhibition, glycoATP remained higher, demonstrating conservation of ISC adaptive response to mitochondrial inhibition across species.

**Figure 3.**
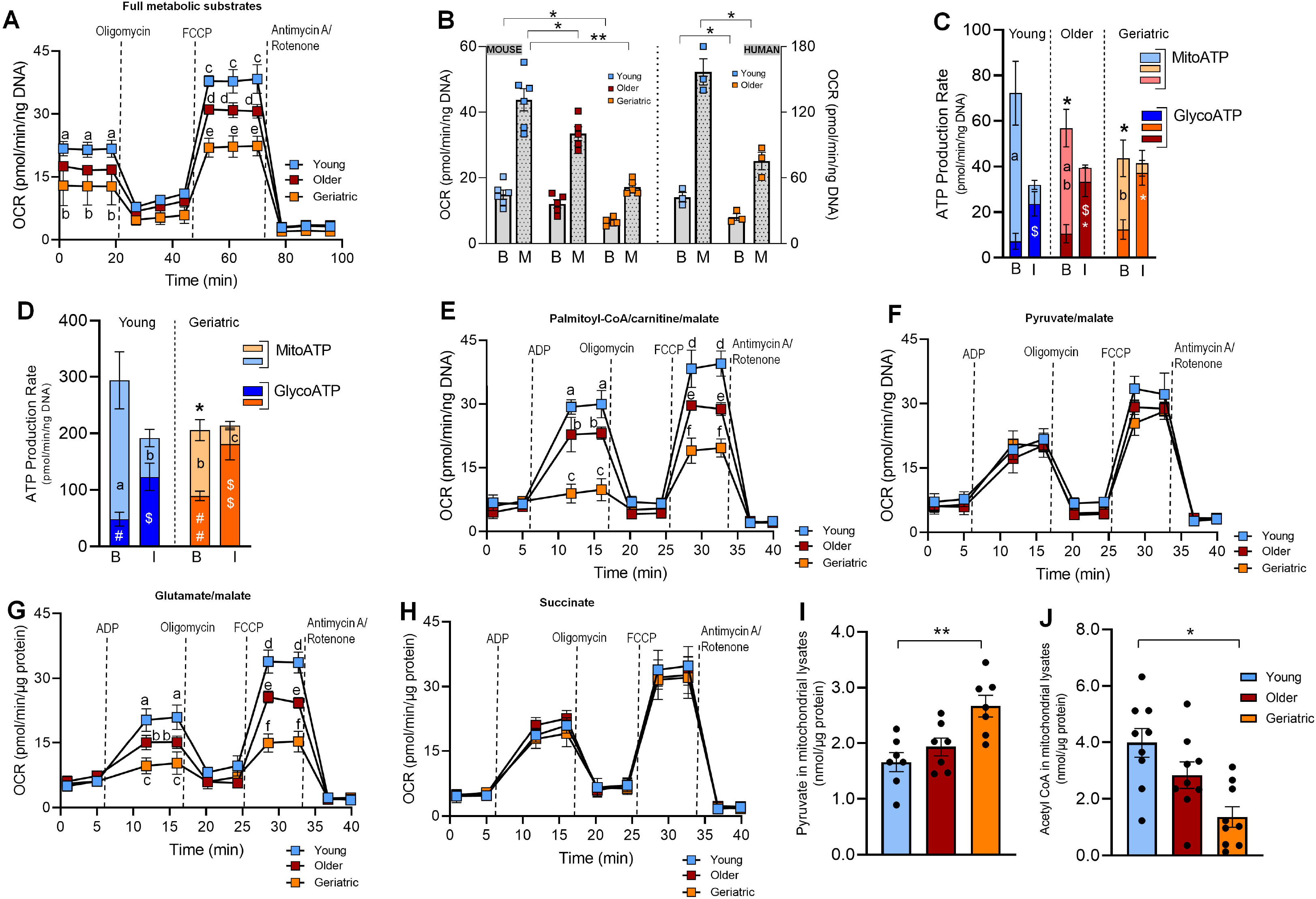
Decreased mitochondrial OCR and mitoATP production in older and geriatric ISCs. **(A)** Mitochondrial OCR profiles in GFP^high^ ISCs sorted from young, older, and geriatric Lgr5-EGFP mice (51-60 enteroids/mouse; *n*=5-6 mice/group) with access to all metabolic substrates in the presence of oligomycin (1.2 μM), FCCP (5 μM), and rotenone (1 μM). Data was analyzed using repeated one-way ANOVA and post hoc Dunn’s test with Benjamini-Hochberg correction. **(B)** Seahorse quantitation of basal (B) and maximal (M) mitochondrial OCR in GFP^high^ ISCs (*left panel*, *n*=5-6 mice/group), and 2-day-old duodenal enteroids generated from young and older individuals (*right panel,* mean of 45-54 enteroids/subject; *n*=3/group), analyzed using Kruskal Wallis ANOVA and post hoc Dunn’s test. **(C)** Total, mito-, and glyco-ATP in GFP^high^ ISCs sorted from Lgr5-EGFP mice (*n*=5-6 mice/group), and 2-day-old duodenal enteroids from young and older individuals **(D,** 45-60 enteroids/subject; *n*=3/group), measured at baseline (B) and after inhibition (I) by oligomycin and antimycin/rotenone. Data were analyzed using two-way ANOVA with post hoc Tukey’s test (mito- and glycol-ATP), or Scheirer–Ray–Hare test and post-hoc Dunn test with Benjamini-Hochberg correction (total ATP). OCR profiles in mitochondria isolated from GFP^high^ ISCs (*n*=5-6 mice/group), and treated with single metabolic substrates, 40 μM palmitoyl-CoA/carnitine **(E)**, 5 mM pyruvate **(F)**, 5 mM glutamate **(G)**, and 10 mM succinate **(H)** in the presence of 5 mM malate. Pyruvate **(I)** and acetyl CoA **(J)** in mitochondria isolated from GFP^high^ ISCs from Lgr5-EGFP mice (*n*=7-9 mice/group), analyzed using non-parametric one-way ANOVA and post hoc Dunn’s test (*, p<0.05; **, p<0.01). Data points with different letters are significantly different (A, E-H). Groups with different letters and symbols indicate significant difference between mito- and glycoATP, respectively (C, D). Asterisks represent significant differences in total ATP compared to the young.

To gain further insights into mechanisms underlying mitochondrial metabolism, we isolated mitochondria from GFP^high^ ISCs and measured respiratory responses to single metabolic substrates. Older and geriatric ISC mitochondria displayed a progressive decline in OCR in the presence of palmitic acid **(Figure 3E, Supplemental Figure 3C)**. Maximal OCR in the geriatric cohort was less than half of that observed in the young **(Figure 3E)**. Similar results were observed with linoleic and linolenic acids (data not shown). In contrast, when pyruvate was used, OCR of older and geriatric ISC mitochondria was comparable to that of young mitochondria **(Figure 3F, Supplemental Figure 3D)**, demonstrating that aging is associated with a shift in preference for pyruvate over FAs. OCR was also suppressed in the presence of glutamate **(Figure 3G, Supplemental Figure 3E)**. Notably, succinate, the substrate for complex II of the electron transport chain (ETC), induced comparable OCRs across all age groups **(Figure 3H),** validating ETC functionality. Taken together, these mechanistic data confirm that aging alters fuel selection of ISCs from FAs to pyruvate.

### Aging mediated decrease in mitochondrial ATP production is due to decreased PDH activity

We then measured pyruvate and acetyl CoA levels in mitochondria isolated from GFP^high^ ISCs. Although pyruvate levels in ISCs enteroids were 60% higher than young ISCs **(Figure 3I)**, acetyl CoA levels were 66% lower **(Figure 3J),** suggesting possible decrease in pyruvate dehydrogenase (PDH) activity. Indeed, PDH activity was suppressed in older (-55%) and geriatric ISCs (-76%), compared to the young **(Figure 4A, B)**. PDH is regulated via phosphorylation of its serine residues by PDH kinases (PDKs) that inhibit activity. Phospho-PDH expressions (Ser 293 and Ser 300) were significantly higher in geriatric ISCs **(Figure 4C, D)**, collectively demonstrating that aging decreases PDH activity in ISCs.

**Figure 4.**
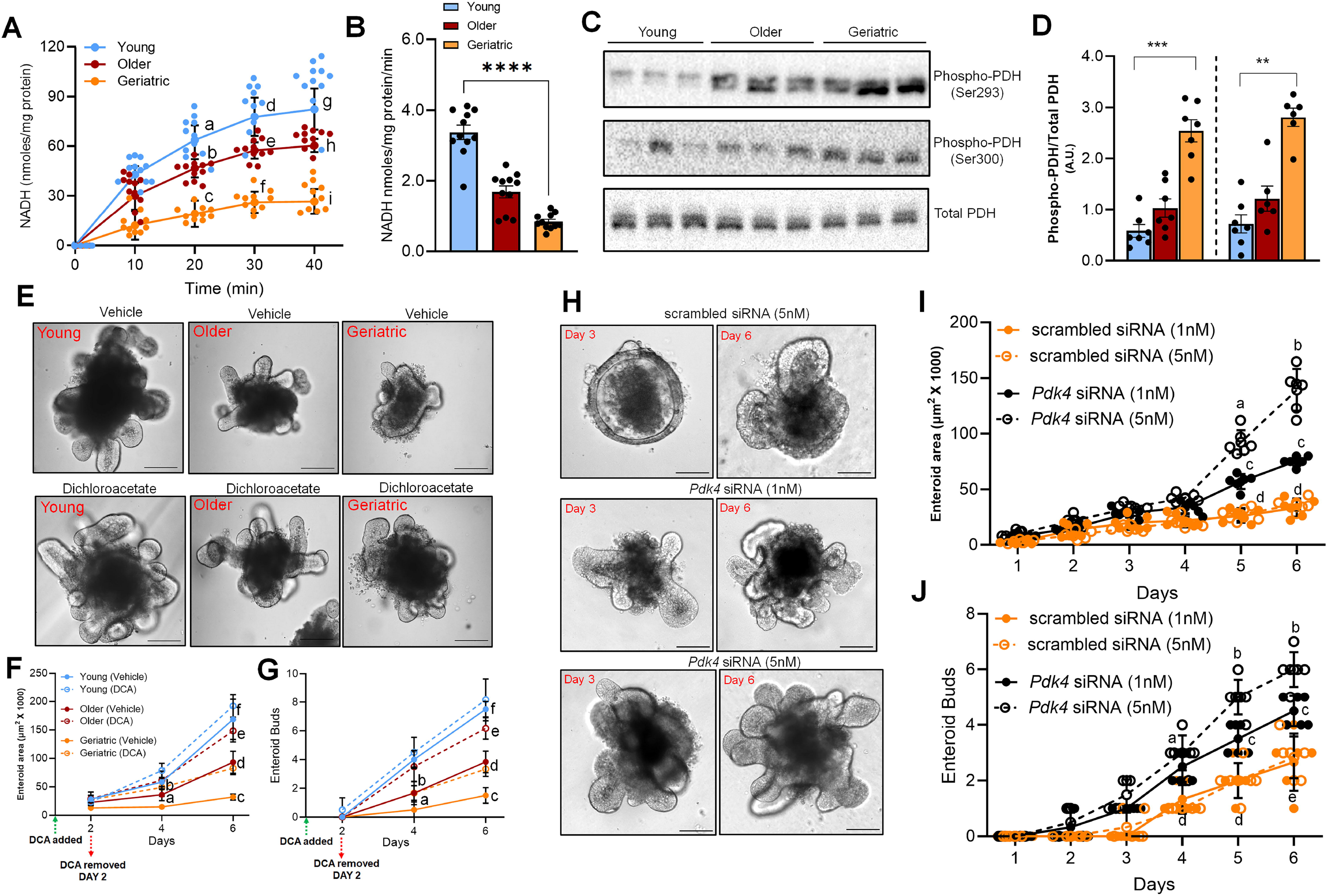
PDK4 inhibition and subsequent PDH upregulation robustly enhances stemness in older and geriatric ISCs. Low PDH activity in GFP^high^ ISCs isolated from young, older, and geriatric Lgr5-EGFP mice (*n*=10-11 mice/group) and grown in culture for 2 days **(A, B)**. Representative blot **(C)**, and quantitation **(D)** of phospho-PDH (Ser293 and Ser300) and total PDH expressions in GFP^high^ ISCs (*n*=6-7 mice/group). Brightfield images **(E)**, area **(F)** and bud numbers **(G)** of enteroids generated from GFP^high^ ISCs isolated from Lgr5-EGFP mice, treated with PBS vehicle or DCA (4mM) for 2 days and imaged after 6 days of seeding. Data was analyzed using Scheirer–Ray–Hare test and post-hoc Dunn test with Geisser-Greenhouse correction for sphericity. Brightfield images **(H)**, area **(I)** and bud numbers **(J)** of enteroids from GFP^high^ ISCs, transfected with scrambled or Pdk4 siRNA (1- and 5 nM) for 2 days and imaged after 3 - 6 days of seeding. Data was analyzed as in 3A (**, p<0.01; ***, p<0.001; ****, p<0.0001). Data points with different letters are significantly different (A, F, G, I, J).

### Suppression of PDK4 enhances PDH activity and robustly improves stemness in older and geriatric enteroids

Given the low PDH activity in aging ISCs, we determined whether enhancing PDH activity improves stemness. PDH is maintained in active, non-phosphorylated form by inactivating pyruvate dehydrogenase kinase (Pdk4) ^8, 12^. We suppressed Pdk4 activity using two approaches: pharmacological inhibition by dichloroacetate (DCA) and transient silencing of *Pdk4* gene using siRNA. DCA (4nM) was added during harvest and treated for 2 days, following which it was removed to minimize effects on enterocytes that also express Pdk4. DCA treatment significantly increased bud numbers and bud area in the older enteroids, an effect that was potentiated in the geriatric ISCs (**Figures 4E-G**). Enteroid area increased by 32-54% and 108-153% (**Figures 4E, F)**, while bud numbers increased by 32-45% and 167-227% in the older and geriatric enteroids, respectively **(Figures 4E, G**). Similar effects were observed after transient silencing of *Pdk4* gene. Scrambled or *Pdk4* siRNA was added during harvest and media was changed after 2 days. Compared to controls, geriatric enteroids transfected with *Pdk4* siRNA showed progressive increase in enteroid area and bud numbers in a dose-dependent manner **(Figure 4H-J)**. Enteroid area increased by 108-203% (**Figures 4H, I)**, while bud numbers increased by 116-227% (**Figure 4H, J)** after transient *Pdk4* silencing. Thus, *Pdk4* suppression robustly enhanced growth and budding capacities of older and geriatric enteroids, making them virtually indistinguishable from the next immediate younger cohort.

### Pdk4 inhibition enhances stemness via increasing mitochondrial utilization of pyruvate, OCR, and mitoATP

We then investigated mechanisms underlying Pdk4 mediated enhancement of stemness. Geriatric GFP^high^ ISCs treated with DCA or transfected with *Pdk4* siRNA showed a ∼2 increase in basal and ∼85-94% increase in maximal OCR, respectively **(Figures 5A, B)**. Similar improvement in stemness was observed in older GFP^high^ ISCs **(Supplemental Figures 4A, B)**. Additionally, *Pdk4* inhibition also led to a 43-82% increase in basal mitoATP production in geriatric **(Figure 5C)** and older ISCs **(Supplemental Figure 4C)**, while slightly lowering glycoATP levels. To further delineate the metabolic substrate preferred for increased mitoATP, we isolated mitochondria from treated or transfected GFP^high^ ISCs and measured OCR in response to single metabolic substrates. In geriatric ISCs, Pdk4 suppression did not alter OCR in the presence of palmitic acid **(Figure 5D, Supplemental Figure 4D)**, and linoleic, oleic, and linolenic acid (data not shown), demonstrating that Pdk4 inhibition does not enhance FA utilization. However, a marked increase in maximal OCR (63-66%) was observed with Pdk4 inhibition in the presence of pyruvate **(Figure 5E, Supplemental Figure 4E).** Glutamate evoked a slight increase in OCR **(Figure 5F, Supplemental Figure 4F)**. Taken together, our data show that in aging ISCs, pyruvate shunts away from mitochondrial aerobic oxidation towards anaerobic glycolysis. Pdk4 inhibition re-routes it towards aerobic oxidation and mitoATP formation, leading to robust improvement in growth and budding capacities. Additionally, Pdk4 inhibition reduced mitochondrial pyruvate levels **(Figure 5G)** while concomitantly increasing acetyl CoA levels in geriatric **(Figure 5H)** and older ISCs **(Supplemental Figures 5A, 5B,** respectively), confirming increased mitochondrial pyruvate oxidation.

**Figure 5.**
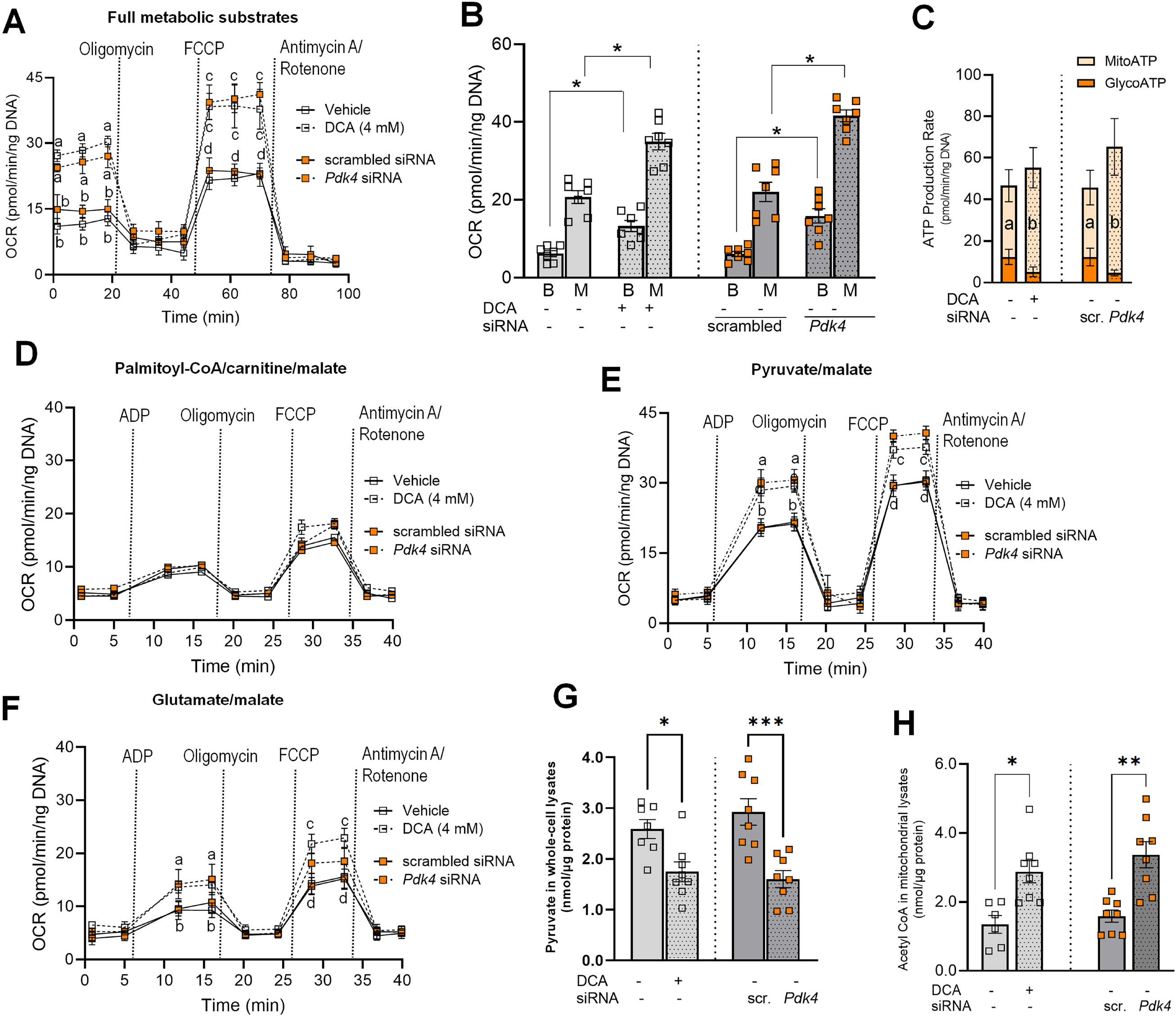
PDK4 inhibition enhances ISC stemness by increasing mitochondrial OCR and mitoATP production. **(A)** Mitochondrial OCR profiles in GFP^high^ ISCs sorted from geriatric Lgr5-EGFP mice treated with vehicle or DCA or transfected with scrambled or Pdk4 siRNA (5 nM) and measured with full access to all metabolic substrates (45-60 enteroids/mouse, *n*=6-7 mice/group/treatment). Data was analyzed as in 4F. **(B)** Seahorse quantitation of basal and maximal mitochondrial OCR in GFP^high^ ISCs from geriatric Lgr5-EGFP mice and analyzed using Kruskal Wallis ANOVA with post hoc Dunn’s test (*n*=6-7 mice/group). **(C)** Total, mito-, and glyco-ATP in GFP^high^ ISCs sorted from geriatric Lgr5-EGFP mice (*n*=6-7 mice/group), treated with vehicle or DCA, or transfected with Pdk4 siRNA. Groups with different letters and symbols indicate significant difference between mito- and glycoATP, respectively. OCR profiles in mitochondria isolated from DCA treated, or transfected GFP^high^ ISCs sorted from geriatric Lgr5-EGFP mice (*n*=6-7 mice/group), after application of single metabolic substrates: palmitoyl-CoA/carnitine **(D)**, pyruvate **(E)**, and glutamate **(F)**. Data was analyzed as described in (A). Pyruvate **(G)** and acetyl CoA **(J)** in mitochondria isolated from GFP^high^ ISCs from geriatric Lgr5-EGFP mice (*n*=7-8 mice), analyzed as in 3J (*, p<0.05; **, p<0.01; ***, p<0.001). For (A) and (D-F), data points with different letters are significantly different.

Similar results were observed in human duodenal enteroids. Both DCA treatment and transient knockdown of *PDK4* robustly enhanced growth and budding capacities of enteroids **(Figures 6A, B).** Enteroid area in the older subjects was 2-3-fold higher after DCA treatment **(Figure 6A, Supplemental Figure 6A)** and transient knockdown (**Figure 6B**, **Supplemental Figure 6A)**, while bud numbers increased by 3-4-fold **(Supplemental Figure 6B**). As observed in mice, PDK4 mediated improvement of stemness in older human enteroids was driven via enhanced mitochondrial OCR. Older enteroids treated with DCA or transfected with PDK4 siRNA showed a 2-fold increase in basal and 75-80% increase in maximal OCR, respectively **(Figure 6C)**, corresponding to a 2-fold increase in basal mitoATP **(Figure 6D).**

**Figure 6.**
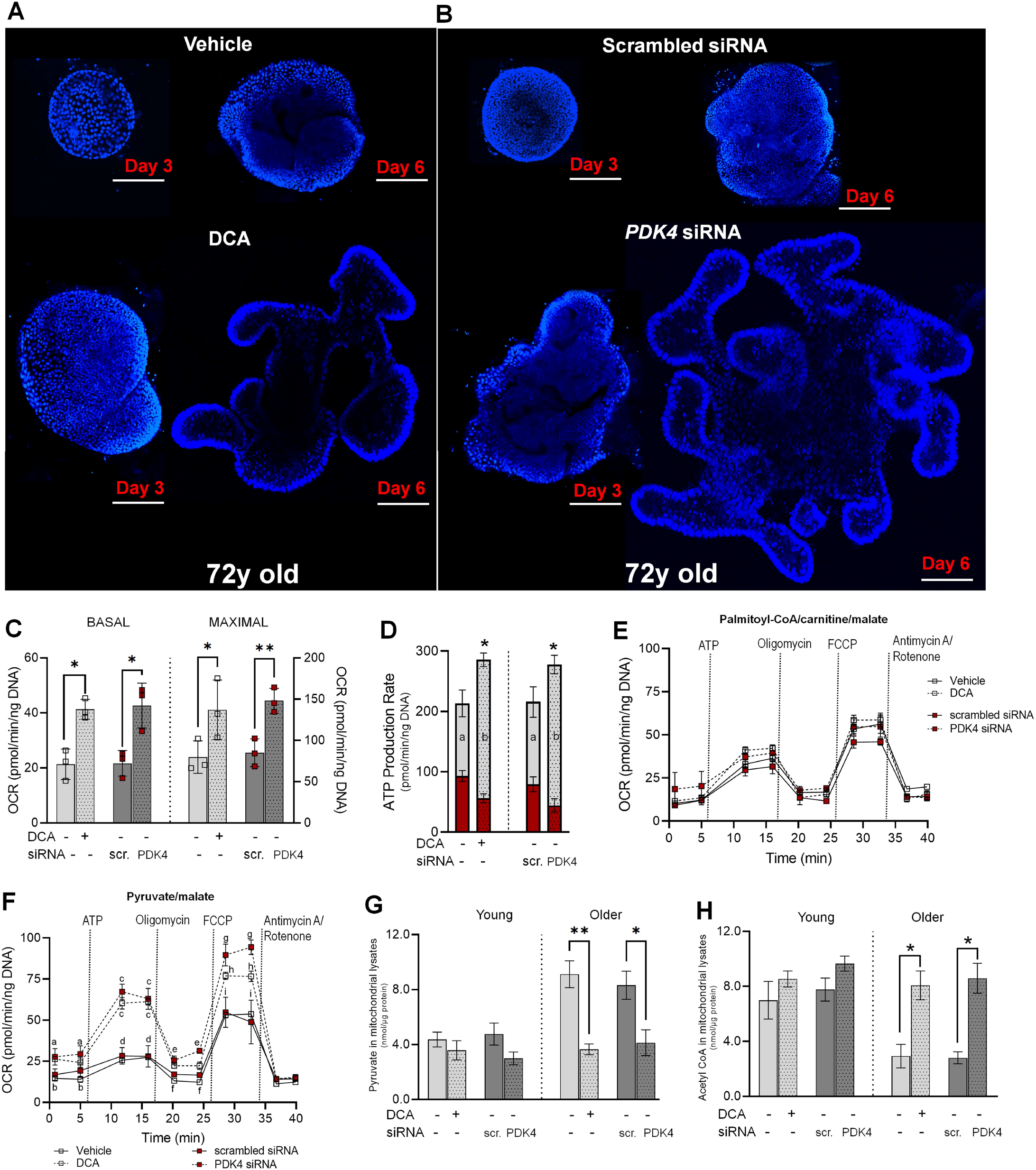
PDK4 inhibition robustly enhances stemness of older human duodenal enteroids via increased OCR and mitoATP. Representative nuclear (DAPI) staining of duodenal enteroids from 72y old female, treated with vehicle or DCA **(A)**, or transfected with scrambled or PDK4 siRNA (10 nM) for 2 days **(B),** and imaged after 3-6 days of seeding. Basal and maximal mitochondrial OCR **(C)**, total, mito, and glycoATP levels **(D)** in duodenal enteroids from 72y old female, treated with vehicle or DCA, or transfected with PDK4 siRNA. Data was analyzed using Scheirer–Ray–Hare test and post-hoc Dunn test with Geisser-Greenhouse correction (*n*=triplicates of 15-20 enteroids each from 3 subjects). Groups with different letters have significantly different mitoATP levels. Asterisks represent differences in total ATP compared to vehicles or scrambled controls. OCR profiles in mitochondria isolated from DCA treated, or transfected enteroids from older subjects, after application of palmitoyl-CoA/carnitine **(E)** and pyruvate **(F)**. Data was analyzed using repeated one-way ANOVA and post hoc Dunn’s test with Benjamini-Hochberg correction (*n*=triplicates of 18-27 enteroids each from 3 subjects). Data points with different letters are significantly different. Pyruvate **(G)** and acetyl CoA **(J)** in mitochondria isolated from enteroids from older subjects, analyzed as in 5D (*, p<0.05; **, p<0.01). Scale bars: (A-B) – 10 µm.

No enhancement in mitochondrial OCR was observed when isolated mitochondria from human enteroids were treated with palmitic acid **(Figure 6E)**, and linoleic, oleic, and linolenic acid (data not shown). However, a marked increase in OCR was observed in the presence of pyruvate **(Figure 6F).** PDK4 downregulation increased maximal OCR by approximately 38% **(Supplemental Figure 6C)**. Glutamate evoked a slight increase in OCR **(Supplemental Figure 6D)**. As observed in mice, mitochondrial pyruvate levels were higher in enteroids from older individuals **(Figure 6G)**. PDK4 suppression significantly lowered pyruvate content to levels comparable to younger enteroids **(Figure 6G)**, while increasing acetyl CoA levels **(Figure 6H)**, demonstrating enhanced pyruvate utilization.

Finally, to confirm enhanced mitochondrial utilization of pyruvate and acetyl CoA with Pdk4 inhibition, we performed stable isotope tracing using ^13^C_6_-glucose, that robustly labels TCA cycle intermediates ^29^. Sorted GFP^high^ISCs, grown as monolayers and transfected with scrambled or *Pdk4* siRNA were incubated with ^13^C_6_-glucose for 6 hours and ^13^C enrichment in TCA cycle intermediates was measured by LC-MS. Glucose-derived pyruvate enters the TCA cycle via two enzymatic reactions: PDH, which decarboxylates pyruvate to acetyl CoA, and pyruvate carboxylase (PC), which carboxylates pyruvate into oxaloacetate. The PDH route delivers two pyruvate-derived carbons to the TCA cycle, while PC delivers three carbons (**Figure 7A**). Hence, citrate (M+2): pyruvate (M+3), and citrate (M+3): pyruvate (M+3) labeling are surrogates for PDH-dependent and PC-dependent entry, respectively. Our data show that citrate (M+2): pyruvate (M+3) ratio exceeded citrate (M+3): pyruvate (M+3) (**Figure 7B**), demonstrating that PDH mediated decarboxylation is the dominant route for pyruvate derived carbon entry in ISCs. The percentage of labeled (M+2) isotopologue of citrate was significantly lower in the geriatric ISCs **(Figure 7C)**. Pdk4 silencing in the geriatric enhanced labeling to levels virtually indistinguishable from younger counterparts **(Figure 7C).** Label enrichment was also observed in other TCA cycle intermediates, including alpha ketoglutarate (Alpha-KG, **Figure 7D),** succinate **(Figure 7E)**, and malate **(Figure 7F)**, demonstrating that aging leads to decreased carbon influx into the TCA cycle from glucose and glucose-derived derived metabolites. Pdk4 inhibition restores influx, subsequently enhancing mitochondrial OCR and mitoATP production.

**Figure 7.**
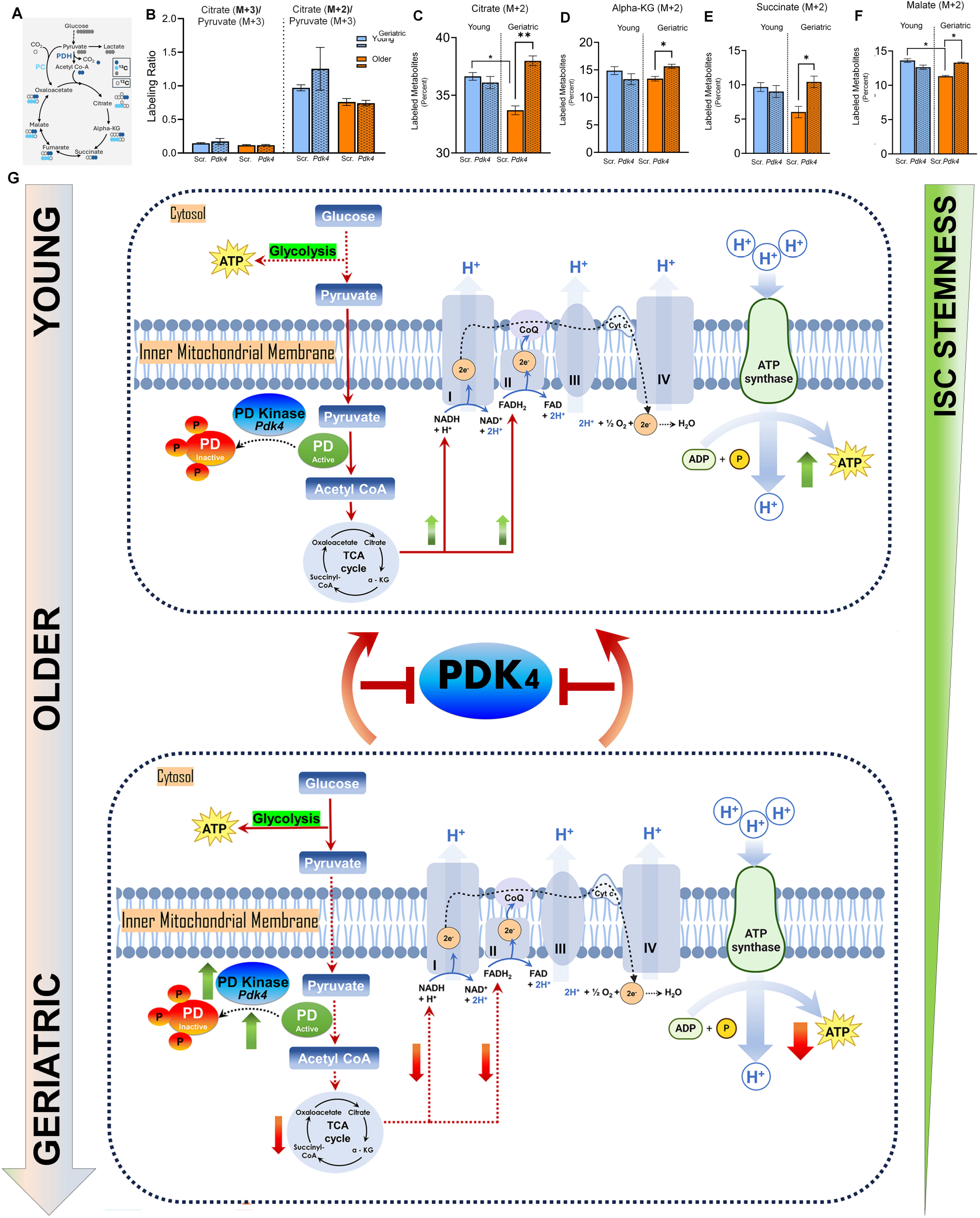
Enrichment of carbohydrate-derived carbon ^13^C_6_ in TCA cycle intermediates upon PDK4 inhibition in geriatric ISCs. **(A)** Metabolic routes for glucose-derived pyruvate in the TCA cycle. **(B)** Citrate (M+2): pyruvate (M+3) and citrate (M+3): pyruvate (M+3) ratio in GFP^high^ ISCs from young and geriatric Lgr5-EGFP mice (45-60 enteroids, *n*=3 mice/group), grown as monolayers for 2 days, and transfected with scrambled or *Pdk4* siRNA (5 nM). **(C-F)** Percentage enrichment of labeled (M+2) isotopologue in citrate **(C)**, alpha-KG **(D)**, succinate **(E)**, and malate **(F)** in young and geriatric Lgr5-EGFP mice (45-60 enteroids, *n*=3 mice/group) and transfected. *Pdk4* siRNA. Data was analyzed using non-parametric one-way ANOVA and post hoc Dunn’s test. **(G)** Schematic of mechanism for PDH: PDK mediated enhancement of ISC stemness.

Collectively, our data documents that aging is associated with marked decrease in OCR and mitoATP production, resulting from decreased PDH activity **(Figure 7G),** and a shift in preference from FA to glucose as fuel source. PDK4 inhibition restores PDH activity and robustly enhances stemness of human and mouse ISCs by driving glucose-derived carbon entry into TCA cycle, subsequently increasing mitochondrial OCR and mitoATP **(Figure 7G),** notably without influencing FA utilization.

## DISCUSSION

Aging impairs regenerative capacity of ISCs ^2, 30^ which affects epithelial homeostasis ^31, 32^. Since the pioneering finding by Potten *et al*., that demonstrated slow epithelial turnover in aged mice ^32^, most studies have focused on the transcriptional landscape regulating ISC niche. Our study reveals novel and significant roles of mitochondrial metabolite handling in rescuing age-induced decline in ISC stemness and function. We demonstrate that PDH driven mitochondrial oxidation of the master metabolic substrate pyruvate robustly enhances ISC stemness in the proximal small intestine. Mechanistic analyses using full and single metabolic substrates on isolated mitochondria showed that stemness is enhanced by increasing mitochondrial OCR, and mitoATP generation. PDH and PDK4 play pivotal roles in driving mitochondrial oxidation of the carbohydrate fuel and subsequently ameliorating age-induced decline in ISC stemness.

The biology of aging manifests itself as a slow process in eukaryotic cells. Our study demonstrating a progressive decline in ISC stemness spanning across the 2-year-old lifespan of a mouse, reveals that age associated decline in stemness begins as early as 8-10 months. However, no significant changes were observed in Lgr5, Olfm4, Asl2, Axin 2, Cd44 expressions in GFP-sorted cells across all ages. These observations concur with previous findings, that noted no significant changes in the ISC molecular signature ^2, 33, 15, 34^, confirming that ISC function, rather than absolute numbers is impacted by aging. Interestingly, decline in stemness was recapitulated in older humans, too. This is the first study, to our knowledge, that reports age-induced decline in growth and budding capacities of human duodenal ISCs. Inability to label human ISCs limits accurate quantitation of ISC numbers in human enteroids, but recapitulation of the enteroid phenotype possibly stems from conservation of molecular signature pattern. Indeed, 2-3-day-old enteroids from young and older individuals showed no significant differences between *LGR5*, *OLFM4*, *ASL2*, *AXIN 2*, *CD44* mRNA expressions (data not shown). Historically, ISC metabolism is recognized as a permissive consequence of the cellular (proliferative or differentiated) state. However, recent studies have demonstrated that nutrient metabolism may impact ISC function and fate ^35, 36^. Dietary FAs were sufficient to recapitulate *in vivo* stem-cell phenotype in an autonomous manner, with minimal niche dependence ^37^. Our study reveals that this phenomenon is age dependent. While addition of palmitic and oleic acid to cultures enhanced enteroid formation ^37^, we observed that blocking mitochondrial FA transport severely blunted growth and budding capacities, most profoundly in younger mice and humans. Geriatric ISCs were virtually unaffected. Moreover, OCR and mitoATP production robustly increased when mitochondria from young ISCs were incubated with FAs, but no significant enhancements were observed in the geriatric or older mitochondria. This is less likely due to differences in mitochondrial biogenesis, as numbers of ISC mitochondria were similar across all age groups. Low CPT1 activity possibly limits mitochondrial FAO transport and primes towards increased glucose utilization. However, given the accumulation of pyruvate and low acetyl CoA, and low PDH activity in aging ISC mitochondria, anaerobic glycolysis is indicated as the preferred route for ATP. The modest elevation in glycoATP in aging human and mice ISCs corroborates this interpretation. ISCs, unlike many stem cells, rely on OXPHOS for ATP ^38^. Our finding that aging ISCs have increased propensity to switch from OXPHOS to glycolysis is significant and may offer relevant insights to observed shift from self-renewal to commitment to the differentiated phenotype with aging. A decline in mitotic cells was observed in the aged crypts ^2^. Fewer molecules of ATP generated via anaerobic fermentation may explain the decline in cells undergoing mitosis. Alternatively, given that ATP generation via this pathway is quicker and requires less machinery, the shift to glycolysis may constitute a possible compensatory mechanism in aging ISCs to cater to energy demands during mitosis.

PDH, known to function as mitochondrial gatekeeper occupies a central node of intermediary metabolism by linking glycolysis to TCA cycle ^8, 12^. Regulation of PDH activity by manipulating post-translational serine phosphorylation is studied extensively in cancer research ^5, 8, 9^. In our study, inhibition of PDK4 (that phosphorylates and inactivates PDH) robustly enhanced stemness of aging ISCs by driving the oxidative decarboxylation of pyruvate to acetyl CoA. Transient (instead of stable) knockdown of the PDK4 gene in conjunction with the 2-day transfection window, minimized confounding effects of enterocytes which retain PDK4 after terminal differentiation. Delivery of siRNA into matrigel-enteroid suspensions during plating and the use of serum significantly improved transfection efficiency (∼78%). Metabolite flux analyses following ^13^C-glucose incubation revealed that PDK4 inhibition augmented enrichment of labeled (M+2) isotopologue in TCA cycle intermediates, demonstrating restoration of mitochondrial oxidation in aging ISCs. We speculate that enhanced OCR and increased ATP production recalibrates ISC fate, priming towards improved self-renewal and viability of parent ISCs. Indeed, mitochondrial regulation was recently observed to enhance self-renewal in neural and hematopoietic stem cells, attributable, in part, to increased NADH pool, cholesterol biosynthesis and extracellular vesicle biogenesis and release ^26, 39^. Future studies are warranted to dissect the impact of PDH upregulation on ISC self-renewal versus differentiation and how mitochondrial OXPHOS governs proliferative commitment in the cell cycle. Recently, Lgr5+ ISCs were reported to spend a significant proportion of the G_1_ phase of the cell cycle in an unlicensed state until proliferative-fate decisions ^40^. Duration of the unlicensed G_1_ state was inferred to be a key regulator of ISC proliferation. We speculate that aging alters the switch between proliferative fate decisions and quiescence and is driven by declining propensity towards mitochondrial OXPHOS.

A body of preclinical *in vitro* and *in vivo* studies has established the efficacy of PDH upregulation in cancer therapy, with demonstrated effectiveness in a variety of tumors ^41^. The ability of this pathway to “unlock” cancer cells from apoptosis resistance has garnered investment in clinical trials. However, exploiting this pathway for ameliorating and/or reversing hallmarks of aging in ISCs has never been explored. Future studies are warranted to investigate effectiveness in accelerating recovery from radiation damage, improving barrier function, and reducing susceptibility to infections and inflammation.

## ACKNOWLEDGEMENT

We thank Midwestern University (MWU) Core Facility and Dr. Ellen Kohlmeir for technical assistance with the Seahorse XFe24 Cellular Metabolic Analyzer. We thank Karyn DiNovo for assistance with the confocal microscope. We are grateful to the BioCryo facility of Northwestern University’s NUANCE Center, which has received support from SHyNE Resource (NSF ECCS-2025633), the IIN, and Northwestern’s MRSEC program (NSF DMR-2308691) for TEM analyses. We thank the University of Chicago DDRCC, Center for Interdisciplinary Study of Inflammatory Intestinal Disorders (NIDDK P30 DK042086) for human duodenal enteroids used in the study. These studies were supported by MWU startup funds, MWU Outsourcing funds, and CCOM Michael Walczak Research Award (SS), and Kenneth A. Suarez Research Fellowship (GW). Excerpts of data were selected twice for oral presentation at the Digestive Disease Week (DDW) Conference (DDW Chicago 2023, DDW Washington DC 2024).

## REFERENCES

1. Britton E, McLaughlin JT. Ageing and the gut. Proc Nutr Soc 2013;72:173–7.

2. Nalapareddy K, Nattamai KJ, Kumar RS, et al. Canonical Wnt Signaling Ameliorates Aging of Intestinal Stem Cells. Cell Rep 2017;18:2608–2621.

3. Lu V, Roy IJ, Teitell MA. Nutrients in the fate of pluripotent stem cells. Cell Metab 2021;33:2108–2121.

4. Puca F, Fedele M, Rasio D, et al. Role of Diet in Stem and Cancer Stem Cells. Int J Mol Sci 2022;23.

5. Patel MS, Roche TE. Molecular biology and biochemistry of pyruvate dehydrogenase complexes. FASEB J 1990;4:3224–33.

6. Perham RN. Domains, motifs, and linkers in 2-oxo acid dehydrogenase multienzyme complexes: a paradigm in the design of a multifunctional protein. Biochemistry 1991;30:8501–12.

7. Reed LJ. A trail of research from lipoic acid to alpha-keto acid dehydrogenase complexes. J Biol Chem 2001;276:38329–36.

8. Patel MS, Nemeria NS, Furey W, et al. The pyruvate dehydrogenase complexes: structure-based function and regulation. J Biol Chem 2014;289:16615–23.

9. Stacpoole PW, McCall CE. The pyruvate dehydrogenase complex: Life’s essential, vulnerable and druggable energy homeostat. Mitochondrion 2023;70:59–102.

10. Gudi R, Bowker-Kinley MM, Kedishvili NY, et al. Diversity of the pyruvate dehydrogenase kinase gene family in humans. J Biol Chem 1995;270:28989–94.

11. Roche TE, Baker JC, Yan X, et al. Distinct regulatory properties of pyruvate dehydrogenase kinase and phosphatase isoforms. Prog Nucleic Acid Res Mol Biol 2001;70:33–75.

12. Harris RA, Bowker-Kinley MM, Huang B, et al. Regulation of the activity of the pyruvate dehydrogenase complex. Adv Enzyme Regul 2002;42:249–59.

13. Huang B, Gudi R, Wu P, et al. Isoenzymes of pyruvate dehydrogenase phosphatase. DNA-derived amino acid sequences, expression, and regulation. J Biol Chem 1998;273:17680–8.

14. Roche TE, Hiromasa Y, Turkan A, et al. Essential roles of lipoyl domains in the activated function and control of pyruvate dehydrogenase kinases and phosphatase isoform 1. Eur J Biochem 2003;270:1050–6.

15. Barker N, van Es JH, Kuipers J, et al. Identification of stem cells in small intestine and colon by marker gene Lgr5. Nature 2007;449:1003–7.

16. Miura S, Suzuki A. Generation of Mouse and Human Organoid-Forming Intestinal Progenitor Cells by Direct Lineage Reprogramming. Cell Stem Cell 2017;21:456–471 e5.

17. Sato T, Clevers H. Growing self-organizing mini-guts from a single intestinal stem cell: mechanism and applications. Science 2013;340:1190–4.

18. Sato T, Vries RG, Snippert HJ, et al. Single Lgr5 stem cells build crypt-villus structures in vitro without a mesenchymal niche. Nature 2009;459:262–5.

19. Morgan RG, Chambers AC, Legge DN, et al. Optimized delivery of siRNA into 3D tumor spheroid cultures in situ. Sci Rep 2018;8:7952.

20. Ludikhuize MC, Meerlo M, Burgering BMT, et al. Protocol to profile the bioenergetics of organoids using Seahorse. STAR Protoc 2021;2:100386.

21. Millard P, Delepine B, Guionnet M, et al. IsoCor: isotope correction for high-resolution MS labeling experiments. Bioinformatics 2019;35:4484–4487.

22. Pino LK, Searle BC, Bollinger JG, et al. The Skyline ecosystem: Informatics for quantitative mass spectrometry proteomics. Mass Spectrom Rev 2020;39:229–244.

23. Idrovo JP, Yang WL, Nicastro J, et al. Stimulation of carnitine palmitoyltransferase 1 improves renal function and attenuates tissue damage after ischemia/reperfusion. J Surg Res 2012;177:157–64.

24. Bieber LL, Abraham T, Helmrath T. A rapid spectrophotometric assay for carnitine palmitoyltransferase. Anal Biochem 1972;50:509–18.

25. Folmes CD, Nelson TJ, Martinez-Fernandez A, et al. Somatic oxidative bioenergetics transitions into pluripotency-dependent glycolysis to facilitate nuclear reprogramming. Cell Metab 2011;14:264–71.

26. Khacho M, Harris R, Slack RS. Mitochondria as central regulators of neural stem cell fate and cognitive function. Nat Rev Neurosci 2019;20:34–48.

27. Prigione A, Fauler B, Lurz R, et al. The senescence-related mitochondrial/oxidative stress pathway is repressed in human induced pluripotent stem cells. Stem Cells 2010;28:721–33.

28. Basei FL, de Castro Ferezin C, Rodrigues de Oliveira AL, et al. Nek4 regulates mitochondrial respiration and morphology. FEBS J 2022;289:3262–3279.

29. Puchalska P, Huang X, Martin SE, et al. Isotope Tracing Untargeted Metabolomics Reveals Macrophage Polarization-State-Specific Metabolic Coordination across Intracellular Compartments. iScience 2018;9:298–313.

30. Rando TA. Stem cells, ageing and the quest for immortality. Nature 2006;441:1080–6.

31. Martin K, Kirkwood TB, Potten CS. Age changes in stem cells of murine small intestinal crypts. Exp Cell Res 1998;241:316–23.

32. Potten CS, Kovacs L, Hamilton E. Continuous labelling studies on mouse skin and intestine. Cell Tissue Kinet 1974;7:271–83.

33. Kozar S, Morrissey E, Nicholson AM, et al. Continuous clonal labeling reveals small numbers of functional stem cells in intestinal crypts and adenomas. Cell Stem Cell 2013;13:626–33.

34. van der Flier LG, Haegebarth A, Stange DE, et al. OLFM4 is a robust marker for stem cells in human intestine and marks a subset of colorectal cancer cells. Gastroenterology 2009;137:15–7.

35. Alonso S, Yilmaz OH. Nutritional Regulation of Intestinal Stem Cells. Annu Rev Nutr 2018;38:273–301.

36. Mah AT, Van Landeghem L, Gavin HE, et al. Impact of diet-induced obesity on intestinal stem cells: hyperproliferation but impaired intrinsic function that requires insulin/IGF1. Endocrinology 2014;155:3302–14.

37. Beyaz S, Mana MD, Roper J, et al. High-fat diet enhances stemness and tumorigenicity of intestinal progenitors. Nature 2016;531:53–8.

38. Shyh-Chang N, Ng HH. The metabolic programming of stem cells. Genes Dev 2017;31:336–346.

39. Bonora M, Morganti C, van Gastel N, et al. A mitochondrial NADPH-cholesterol axis regulates extracellular vesicle biogenesis to support hematopoietic stem cell fate. Cell Stem Cell 2024;31:359–377 e10.

40. Carroll TD, Newton IP, Chen Y, et al. Lgr5(+) intestinal stem cells reside in an unlicensed G(1) phase. J Cell Biol 2018;217:1667–1685.

41. Michelakis ED, Webster L, Mackey JR. Dichloroacetate (DCA) as a potential metabolic-targeting therapy for cancer. Br J Cancer 2008;99:989–94.

