## Supplementary figures and images for "Mitochondrial oxidation of the carbohydrate fuel driven by pyruvate dehydrogenase robustly enhances stemness of older and geriatric Intestinal Stem Cells"

### Supplemental Figure 1

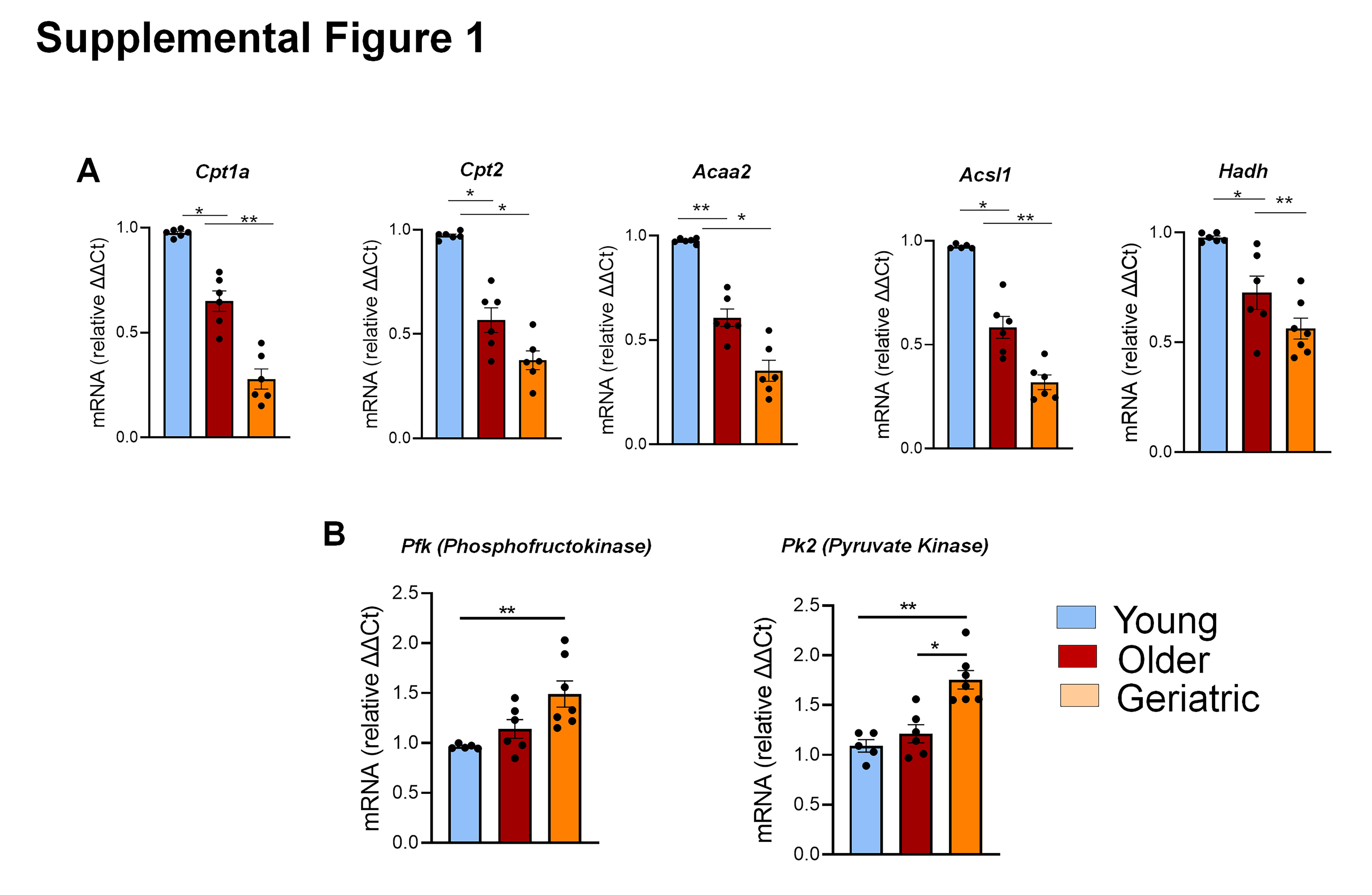

### Supplemental Figure 2

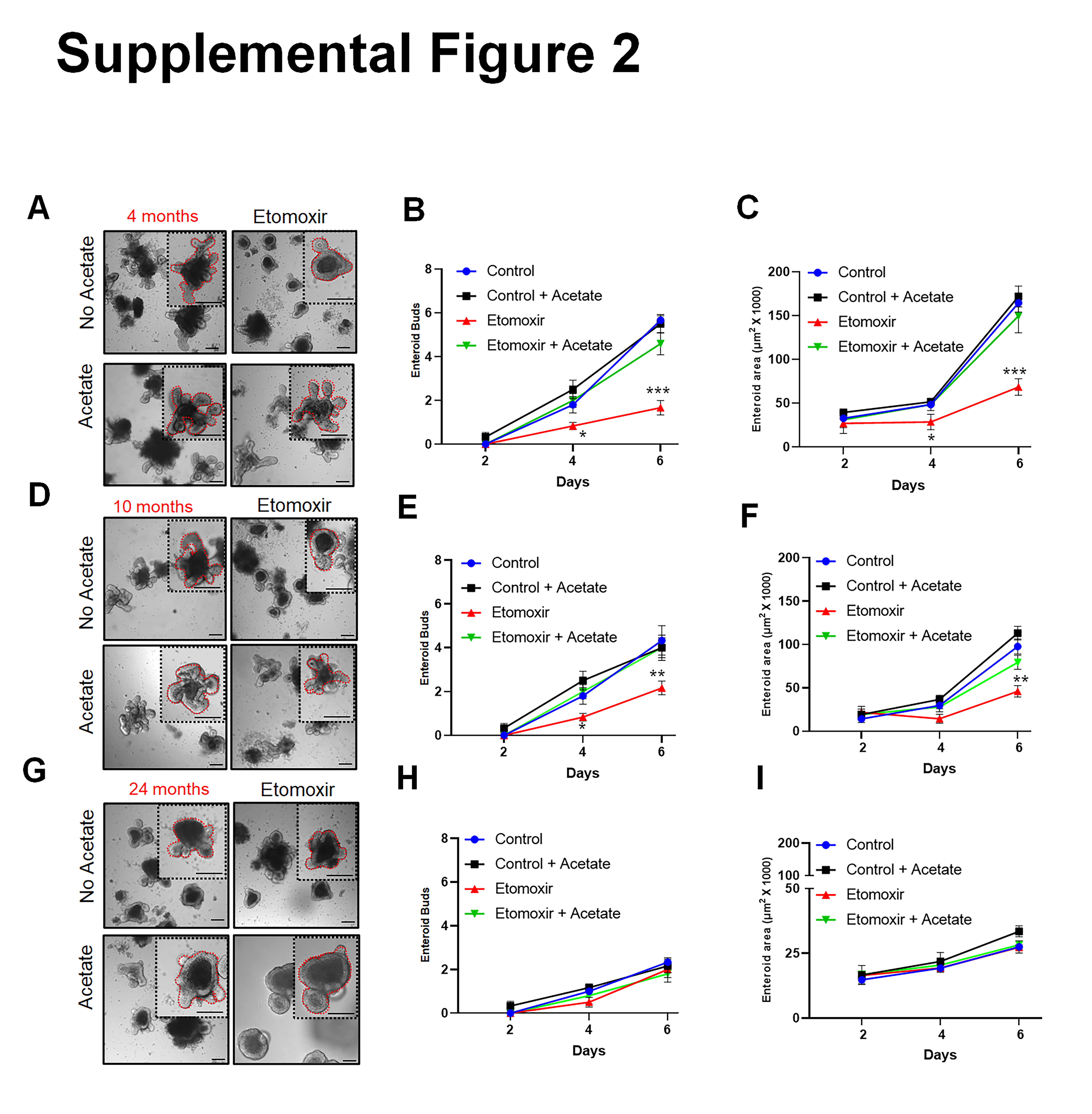

### Supplemental Figure 3

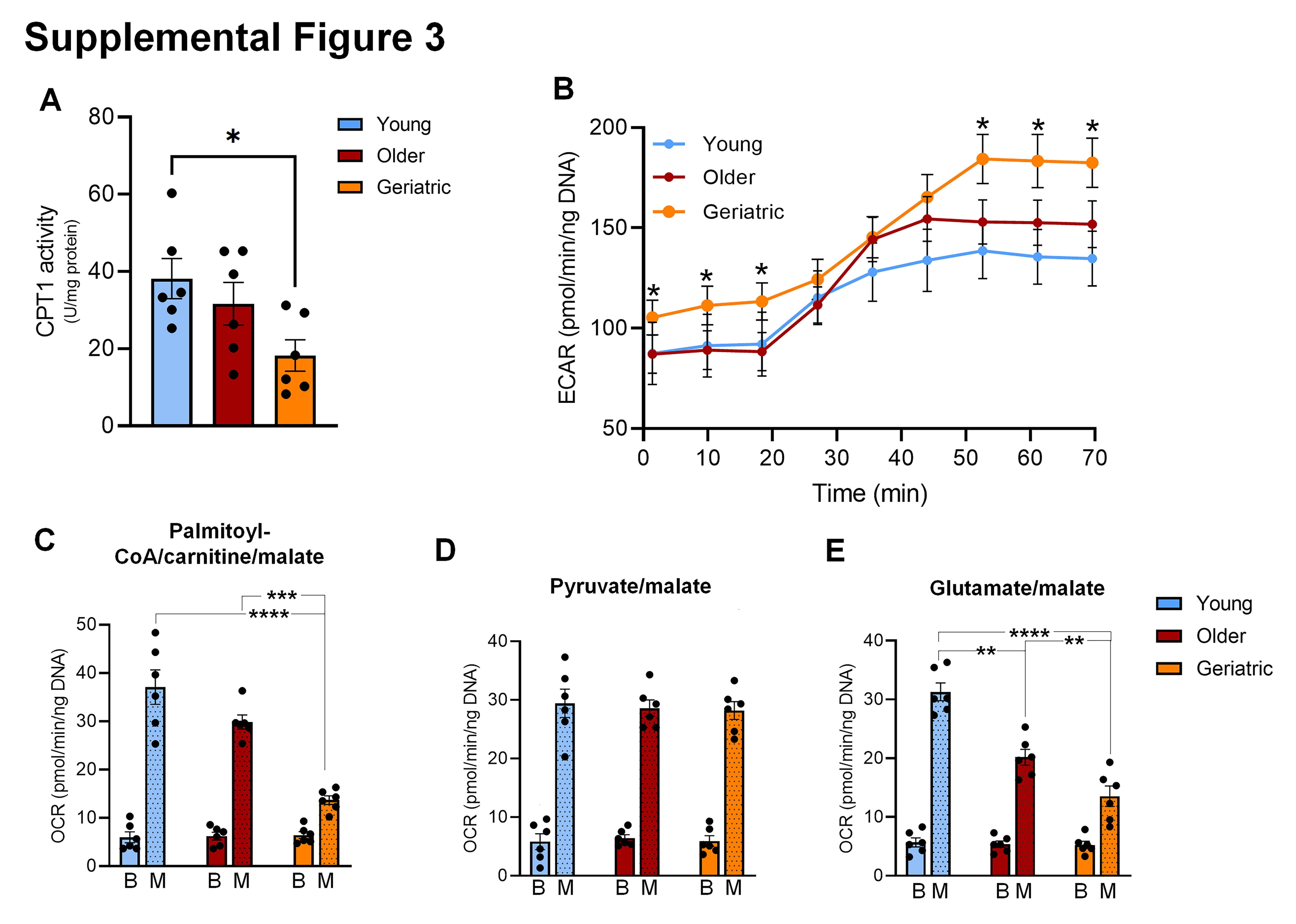

### Supplemental Figure 4

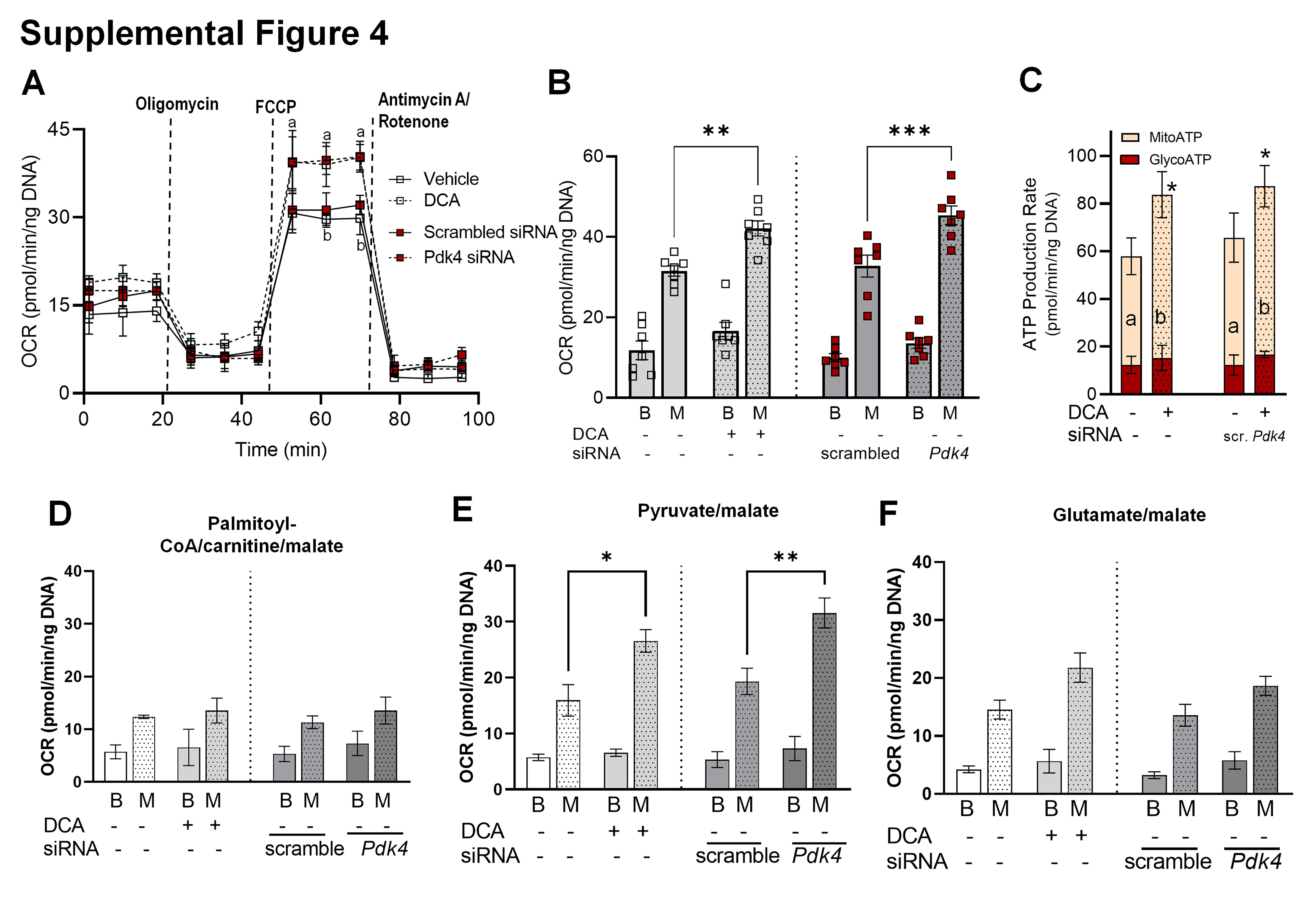

### Supplemental Figure 5

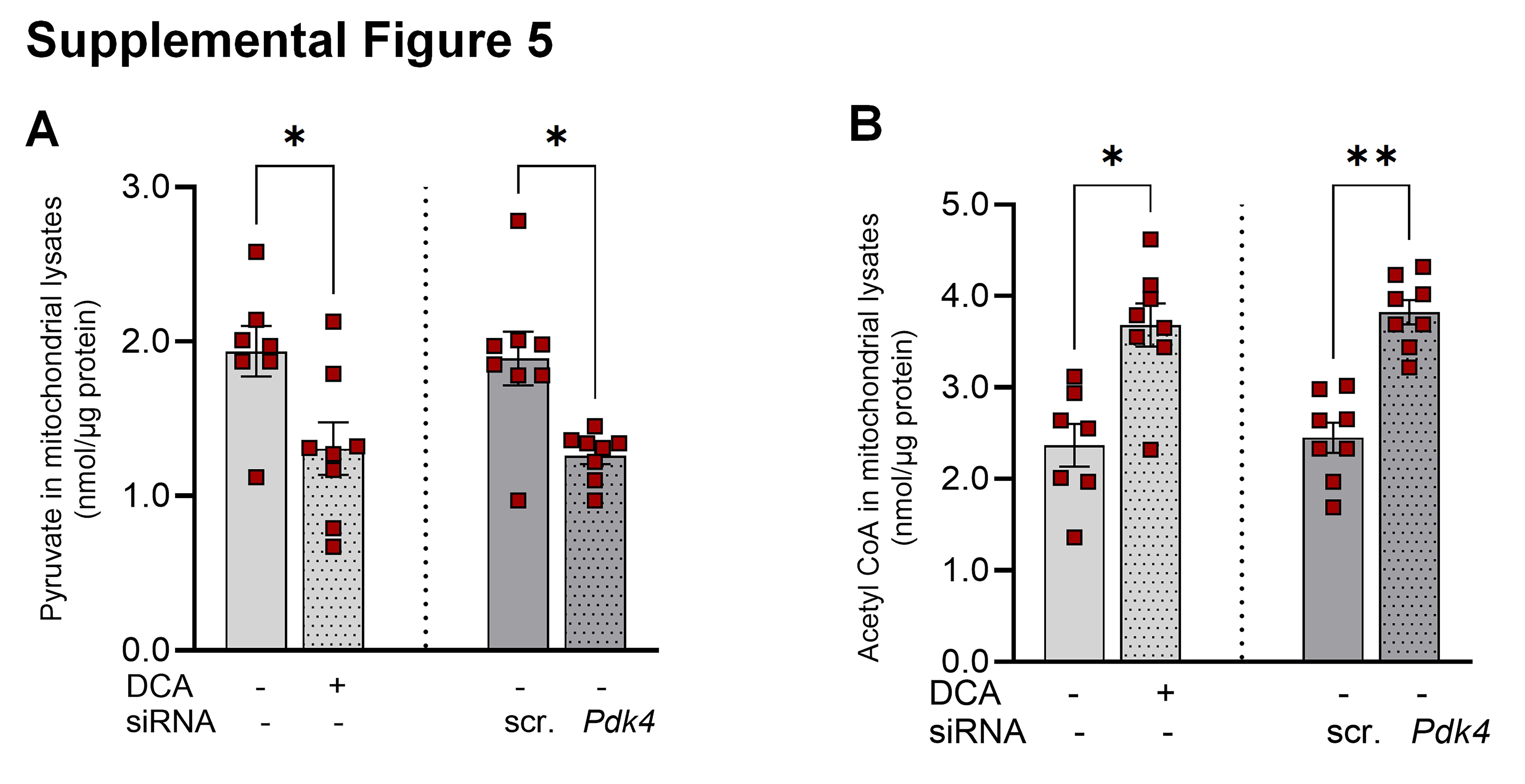

### Supplemental Figure 6

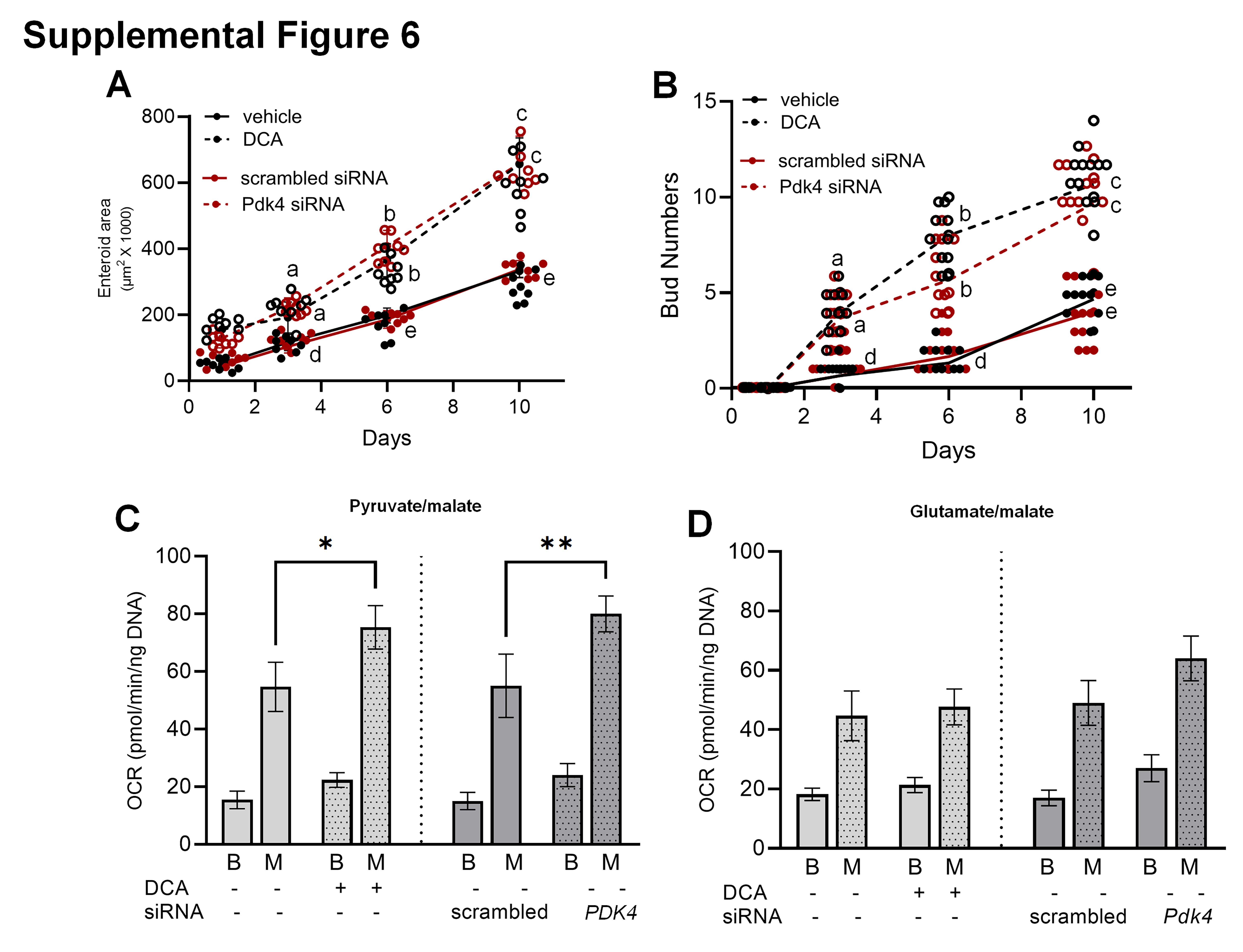
