## Supplemental Information for "Mitochondrial oxidation of the carbohydrate fuel driven by pyruvate dehydrogenase robustly enhances stemness of older and geriatric Intestinal Stem Cells"

**Supplemental Figure Legends**

**Supplemental Figure 1: (A)** *Cpt1a, Cpt2, Acaa2, Acsl1, Hadh, and* **(B)** *Pfk, Pk2* mRNA transcript expressions in GFP-sorted cells (GFP^high^) from proximal small intestinal crypts of young (2-4-months), older (8-10 months), and geriatric (18-24 months) Lgr5-EGFP mice and grown in culture for 2-5 days. Data was normalized to housekeeping transcript *Gapdh* and expressed relative to the young control group. Data was log-transformed and analyzed using repeated one-way ANOVA and post hoc Dunn’s test with Benjamini-Hochberg correction (*n*=mean of triplicates from 7-9 mice/age group). All data are Means ± SEM (*, p<0.05; **, p<0.01).

**Supplemental Figure 2:** Brightfield images **(A)**, bud numbers **(B)**, and enteroid area **(C)** of enteroids generated from single GFP^high^ ISCs isolated from 4-month-old **(A-C),** 10-month-old **(D-F)**, and 24-month-old Lgr5-EGFP mice **(G-I)**, treated with vehicle or etomoxir (50 µM), with or without acetate (1 mM). Enteroid area and bud numbers were analyzed using linear mixed effects modeling with likelihood ratio tests. Graphs depict mean of 3-4 enteroids/passage/mouse (*n*=5-7 mice/age group, 2-3 passages/mouse). Data was analyzed using non-parametric repeated one-way ANOVA and post hoc Dunn’s test with Benjamini-Hochberg correction for multiple comparisons. Asterisks denote significance between groups at the given time point. All data are Means ± SEM (**, p<0.01; ***, p<0.001). Scale bars: (A, D, G) – 20 µm.

**Supplemental Figure 3: (A)** CPT1 activity (U/mg protein) in GFP^high^ ISCs isolated from young (2-4-months)**,** older (8-10-months), and geriatric (18-24 months) Lgr5-EGFP mice. **(B)** Seahorse analyses of extracellular acidification rate (ECAR) in GFP^high^ ISCs isolated from young (2-4-months)**,** older (8-10-months), and geriatric (18-24 months) Lgr5-EGFP mice. **(C-E)** OCR profiles in mitochondria isolated from GFP^high^ ISCs sorted from young, older, and geriatric Lgr5-EGFP mice (*n*=5-6 mice/group), and treated with single metabolic substrates, 40 μM palmitoyl-CoA/40 μM carnitine **(C)**, 5 mM pyruvate **(D)**, and 5 mM glutamate **(E)** in the presence of 5 mM malate, oligomycin (1.2 μM), FCCP (5 μM), and rotenone (1 μM). About 200-300 freshly isolated mitochondria (~10 µg) was used. Data was analyzed using non-parametric repeated one-way ANOVA and post hoc Dunn’s test with Benjamini-Hochberg correction.

**Supplemental Figure 4: (A)** Mitochondrial OCR profiles in GFP^high^ ISCs sorted from older (8-10 months old) Lgr5-EGFP mice treated with vehicle or DCA (4mM) or transfected with scrambled or Pdk4 siRNA (5 nM), and measured with full access to all metabolic substrates in the presence of oligomycin (1.2 μM), FCCP (5 μM), and rotenone (1 μM). About 40-60 enteroids plated across 3 wells were used per mouse. Data was analyzed using non-parametric repeated one-way ANOVA and post hoc Dunn’s test with Geisser-Greenhouse correction (*n*=6-7 mice/group/treatment). Data points with different letters are significantly different. **(B)** Seahorse quantitation of basal (B) and maximal (M) mitochondrial OCR in GFP^high^ ISCs from older Lgr5-EGFP mice and analyzed using non-parametric Kruskal Wallis ANOVA with post hoc Dunn’s test (*n*=6-7 mice/group). **(C)** Total, mito-, and glyco-ATP production rate of GFP^high^ ISCs sorted from older Lgr5-EGFP mice (*n*=6-7 mice/group)**,** treated with vehicle or DCA (4mM) or transfected with scrambled or Pdk4 siRNA (5 nM). Groups with different letters and symbols indicate significant difference between mito- and glycoATP, respectively. Asterisks represent difference in total ATP levels compared to vehicle treated or scramble transfected controls. **(D-F)** OCR profiles in mitochondria isolated from GFP^high^ ISCs sorted from older Lgr5-EGFP mice (*n*=6-7 mice/group), and treated with single metabolic substrates, 40 μM palmitoyl-CoA/40 μM carnitine **(D)**, 5 mM pyruvate **(E)**, and 5 mM glutamate **(F)** in the presence of 5 mM malate, oligomycin (1.2 μM), FCCP (5 μM), and rotenone (1 μM). About 200-300 freshly isolated mitochondria (~10 µg) was used. Data was analyzed as described in (A). All data are Means ± SEM (*, p<0.05; **, p<0.01).

**Supplemental Figure 5: (A)** Pyruvate **(A)** and acetyl CoA content **(B)** in GFP^high^ ISC mitochondria isolated from older Lgr5-EGFP mice (*n*=7-8 mice/group), treated with vehicle or DCA (4mM) or transfected with scrambled or Pdk4 siRNA (5 nM). Data was analyzed using non-parametric one-way ANOVA and post hoc Dunn’s test. All data are Means ± SEM (*, p<0.05; **, p<0.01).

**Supplemental Figure 6: (A, B)** Longitudinal analyses of enteroid area **(A)** and bud numbers **(B)** determined by linear mixed effects modeling with likelihood ratio tests. Graphs depict mean of 3-4 enteroids/passage/subject (*n*=3 subjects/age group, 2-3 passages/subject). Data was analyzed using non-parametric repeated one-way ANOVA and post hoc Dunn’s test with Benjamini-Hochberg correction for multiple comparisons. Data points with different letters are significantly different across the given time point. **(C, D)** Basal (B) and maximal (M) OCR quantitation in mitochondria isolated from DCA (or vehicle) treated, or transfected enteroids from older subjects (*n*=3), after application of single metabolic substrates: 40 μM palmitoyl-CoA/40 μM carnitine **(C)** and 5 mM pyruvate **(D)** in the presence of 5 mM malate, oligomycin, FCCP, and rotenone. About 200-300 freshly isolated mitochondria (~10 µg) was used. Data was analyzed using non-parametric repeated one-way ANOVA and post hoc Dunn’s test with Benjamini-Hochberg correction (*n*=triplicates of 15-25 enteroids each from 3 subjects). All data are Means ± SEM (*, p<0.05; **, p<0.01).

**Supplemental Methods**

**Human and mouse enteroid culture**

Pinch biopsy specimens from human subjects were washed vigorously with ice-cold PBS and incubated in 0.5M EDTA (4°C, 30 minutes). Specimens were then sheared by pipetting, filtered (100µm), and centrifuged (400*g*, 4^°^C, 5 min). Pellets were resuspended in complete growth medium [Advanced DMEM/F12 supplemented with 2mM glutamine, 10 mM HEPES, Wnt conditioned media, N-2 supplement (1X), B-27 supplement (1X), 1.25 mM N-acetylcysteine, 10mM nicotinamide, 50 ng/mL murine EGF, 1µM Jagged 1, 10µM Y-27632, 0.1 ng/mL R-Spondin, 30µM SB202190, 2.5µM Chir99021, 10nM [Leu15]-Gastrin I, 0.5µM LY2157299, 0.5µM A-8301] and plated in Matrigel.

Crypt enriched fractions from the proximal small intestines (16-20 cm distal to the pyloric sphincter) of mice were plated in complete growth medium (Advanced DMEM/F12 with 2mM glutamine, 10 mM HEPES, Wnt conditioned media, N-2 supplement, B-27, 1.25 mM N-acetylcysteine, 10mM nicotinamide, 50 ng/mL murine EGF, 1µM Jagged 1, 10µM Y-27632, and 0.1 ng/mL R-Spondin).

**Western Blot**

Duodenal enteroids or GFP^high^ sorted mouse ISCs were lysed (20 mM Tris, 150 mM NaCl, 1% Triton X-100, 1 mM EDTA, 0.5 mM EGTA). About 30-50 µg of protein was resolved on pre-cast gels and transferred to polyvinylidene difluoride membranes. Blots were blocked in 5% BSA (60 min, RT) and incubated in primary antibodies (Supplementary Table 1) followed by secondary antibodies conjugated to HRP (1:10,000-20,000). Bands were visualized by ECL plus (PerkinElmer) and Chemidoc (BioRad).

**RT-qPCR**

Total RNA was isolated from enteroids using the RNeasy Mini kit (Qiagen). cDNAs were synthesized the iScript cDNA synthesis kit (Bio-Rad, Hercules, CA). Quantitative PCR (qPCRs) was performed on thermal cycler (Bio-Rad) with Platinum Taq DNA polymerase (Invitrogen), SYBR Green dye (Molecular Probes, Carlsbad, CA), and appropriate primers (Supplementary Table 2).

**Statistical Analyses**

Statistical analyses were performed using R (v4.1.2) operating in R Studio. Data was assessed for normality and homoscedasticity using Shapiro–Wilk and Levene’s tests, respectively. Heteroscedastic data sets were log-transformed and re-tested for normality. Normal data were analyzed using one-way ANOVA and Dunn’s post-hoc analyses. Data with multiple repeated measures were analyzed by linear mixed effects modeling with likelihood ratio tests. Repeated measures (one-and two-way ANOVA) and non-parametric data were analyzed using Scheirer–Ray–Hare tests, with Dunn post-hoc analyses with Benjamini-Hochberg correction for multiple comparisons. p < 0.05 was considered significant. Please refer to figure legends for tests used for respective data sets. All data are Means ± SEM and p<0.05 was considered significant.
