## Supplemental Table 1 for "Mitochondrial oxidation of the carbohydrate fuel driven by pyruvate dehydrogenase robustly enhances stemness of older and geriatric Intestinal Stem Cells"

| **Gene** | **Forward** | **Reverse** |
| --- | --- | --- |
| *Acaa2* | CCT CAG TTC TTG TCT GTT CAG | AGG TGT GCG GTG ATT CTG |
| *Acsl1* | TGC CAG AGC TGA TTG ACA TTC | GGC ATA CCA GAA GGT GGT GAG |
| *Cpt1a* | CTC CGC CTG AGC CAT GAA G | CAC CAG TGA TGA TGC CAT TCT |
| *Cpt2* | GCT CCG AGG CAT TTG TC | CAT CGC TGC TTC TTT GGT |
| *Hadh* | AGG CTA CAC GAG CGA GGC GA | ACG GAC CCA TGG GAT ACC CAG C |
| *Pfk* | GGA GAT GCC CAA GGT ATG AAT | GGC GGA CAC TCA GGA ATA AA |
| *Pk 2* | GTG GGT GGA CTA CCA CAA TATC | CAT CAA ACT TCT TCA CGC CTT C |
| *Lgr5* | CGT AGG CAA CCC TTC TCT TAT C | GGA TCA GCC AGC TAC CAA ATA G |
| *Gapdh* | ATG GGA AGC TTG TCA TCA ACG | GGC AGT GAT GGC ATG GAC TG |
| *Olfm4* | AGT GAG GGC AGG TGT ATC T | CTA GAT GCT GCT TAT GGC TCT C |
| *Ascl2* | TAG AGT ACA TTC GGA CCC TCT C | CTC CAC CTT ACT CAG CTT CTT G |
| *c-Myc* | GGG TGG AAG TGA AGC GTT AT | GAC GAG GGA GTT TAG GGA TTT G |
| *CD44* | TAG CAT CTT TGG TGG TGG TG | GTA CCT AGT GTA CAA TGG CTT CTA A |

**Supplemental Table 1: Primers**
