## Supplemental Table 2 for "Mitochondrial oxidation of the carbohydrate fuel driven by pyruvate dehydrogenase robustly enhances stemness of older and geriatric Intestinal Stem Cells"

| **Protein** | **Source** | **Dilution** |
| --- | --- | --- |
| Total Pyruvate Dehydrogenase | Proteintech | 1:8000 |
| Phospho-Pyruvate Dehydrogenase (Ser293) | Proteintech | 1:3000 |
| Phospho-Pyruvate Dehydrogenase (Ser300) | Proteintech | 1:4000 |
| Beta-actin | Cell Signaling Technology | 1:500 |

**Supplemental Table 2: Antibodies**
